# Wasl is crucial to maintain microglial core activities during glioblastoma initiation stages

**DOI:** 10.1101/2021.04.20.440597

**Authors:** Julie Mazzolini, Sigrid Le Clerc, Gregoire Morisse, Cédric Coulonges, Jean-François Zagury, Dirk Sieger

## Abstract

Microglia actively promote the growth of high-grade gliomas. Within the glioma microenvironment an activated (amoeboid) microglial morphology has been observed, however the underlying causes and the related impact on microglia functions and their tumour promoting activities is unclear. Using the advantages of the larval zebrafish model, we demonstrate that pre-neoplastic glioma cells have an immediate impact on microglial morphology and functions. Overexpression of human HRasV12 in proliferating domains of the larval brain induces an amoeboid morphology of microglia, increases microglial numbers and decreases their motility and phagocytic activity. RNA sequencing analysis revealed lower expression levels of the actin nucleation promoting factor *wasla* in microglia. Importantly, a microglia specific rescue of *wasla* expression restores microglial morphology and functions. This results in increased phagocytosis of pre-neoplastic cells and slows down tumour progression. In conclusion, we identified a mechanism that de-activates core microglial functions within the emerging glioma microenvironment.

## Introduction

High grade gliomas represent a complex and devastating disease and are posing an unmet clinical need. These tumours resist multi-modal therapies and survival times are only 14 months on average (Gregory et al., 2020; Kadiyala et al., 2019; Lucki et al., 2019; Wen and Kesari, 2008). In recent years a lot of focus has been on the complex microenvironment of gliomas. Microglia and infiltrating macrophages are the most prominent cell types within the glioma microenvironment and can account for up to 30-50% of the total tumour mass (for review see (Hambardzumyan et al., 2015; Quail and Joyce, 2017). Instructed by a variety of chemokines and cytokines microglia actively promote tumour growth by affecting processes such as cell proliferation and invasiveness, extracellular matrix modifications, angiogenesis and the formation of an immunosuppressive environment (Ellert-Miklaszewska et al., 2013; Hambardzumyan et al., 2015; Komohara et al., 2008; Markovic et al., 2005; Pyonteck et al., 2013; Wang et al., 2013; Wu et al., 2010; Zhai et al., 2011; Zhang et al., 2012). While these processes have been described in gliomas, surprisingly little is known about the apparent change of morphology of microglia within the glioma and the possible impact on their functions. Microglia, as the resident innate immune cells of the brain, display unique morphological features. Under physiological conditions microglia are in a surveillance mode and actively and continuously scan their microenvironment using dynamic large processes providing them a ramified morphology (Nimmerjahn et al., 2005). However, once the homeostasis is altered by injury or brain pathologies, microglia retract their processes to acquire an amoeboid shape and become “activated”. This activated state can correlate with an either anti- or pro-inflammatory state of microglia (Bernier et al., 2019; Bolasco et al., 2018; Chia et al., 2018; Karperien, 2013; Kettenmann et al., 2011; Lawson et al., 1992; Madry et al., 2018a, 2018b). Of note, an activated (amoeboid) microglial morphology has been observed *in-vivo* across different glioma models at different stages of glioma growth as well as within human glioma samples (Annovazzi et al., 2018; Chia et al., 2018; Juliano et al., 2018; Kvisten et al., 2019; Resende et al., 2016; Ricard et al., 2016). Furthermore, these activated microglia show a decreased phagocytic activity and motility within the central area of neoplastic lesions (Hutter et al., 2019; Jaiswal et al., 2009; Juliano et al., 2018; Pyonteck et al., 2013; Wu et al., 2010). The mechanisms underlying this rapid and drastic morphological remodelling are still not known. Clearly, these morphological phenotypes must be highly regulated and involve adaptations to the cellular cytoskeleton (Bernier et al., 2019; Okazaki et al., 2020). Cell morphology, phagocytosis and motility are cellular processes known to be actin-dependent and are crucial for the multitasking roles of microglia (Koizumi et al., 2007; Liu et al., 2020; Lively and Schlichter, 2013; Okazaki et al., 2020; Pollard and Cooper, 2009; Uhlemann et al., 2015). An alteration of the microglial actin cytoskeleton and their morphology will most likely affect several of these core functions. Therefore, it is important to understand the impact of an activated microglial morphology on their functions within the tumour microenvironment and its effects on tumour growth.

Here, we investigated the influence of a pre-neoplastic glioma environment on microglia morphology and related functions. We utilised a recently published zebrafish glioblastoma multiforme (GBM) model which is based on expression of the human oncogene HRasV12 in the proliferating domains of the developing brain and gives rise to tumours similar to the mesenchymal subtype of human GBM (Mayrhofer et al., 2017). Analysing larval stages of this zebrafish model allowed us to directly study the influence of an early pre-neoplastic environment on the morphology and functions of microglia *in vivo*. Importantly, we detected an immediate impact of pre-neoplastic HRasV12^+^ cells on the microglia population resulting in an activated phenotype and increased proliferation of the microglia. Furthermore, their phagocytic activity, motility and speed was significantly reduced compared to control microglia. RNA sequencing of microglia revealed significantly lower expression levels of *wasla*, the zebrafish orthologue of human WASP like actin nucleation promotion factor (WASL, also known as N-WASP), a key regulator of actin cytoskeleton organization (Dart et al., 2012; Linder et al., 1999; Lorenz et al., 2004; Park and Cox, 2009; Yamaguchi et al., 2005; Yu et al., 2012). lmportantly, a microglia specific rescue of *wasla* expression in HRasV12^+^ larvae restored microglial morphology as well as their number, speed and motility. Furthermore, the *wasla* rescue in microglia restored their phagocytic activity which resulted in improvements in both engulfment of pre-neoplastic cells and survival.

## Results

### Pre-neoplastic cells affect the microglia population in the larval zebrafish brain

Although an activated morphology is a consistent feature of microglia within gliomas and has been described across models and species (Chia et al., 2018; Hutter et al., 2019; Kvisten et al., 2019; Ricard et al., 2016), the underlying causes and the timing of the change in morphology are not understood.

Here, we investigated the effect of pre-neoplastic cells on the microglia population using a zebrafish GBM model (Mayrhofer et al., 2017). This model is based on overexpression of human HRasV12 in the proliferating regions of the developing central nervous system (CNS) and results in aggressive tumours resembling human mesenchymal GBM (Mayrhofer et al., 2017). To analyse pre-neoplastic stages, we outcrossed the driver fish line *Et(zic4:GAL4TA4,UAS:mCherry)_hmz5_* (Distel et al., 2009) to the line *Tg(UAS:eGFP-HRasV12)_io006_* (Santoriello et al., 2010) (hereafter Zic/HRasV12 model) (Fig1A). The Zic4 enhancer drives Gal4 expression in the proliferating regions of the developing central nervous system (CNS) and upon binding of Gal4 to its target UAS sequence, activates expression of mCherry in control larvae (hereafter HRasV12^-^) and additional eGFP-HRasV12 expression in tumour developing fish (hereafter HRasV12^+^ larvae). In this model, HRasV12 expression is enriched in the telencephalon from 1 day post fertilisation (dpf) onwards, followed by the tectal proliferation zone (TPZ) at 2 dpf and in the cerebellum from 3 dpf. The expression is low in the optic tectum (Fig1A). This set up results in significantly increased proliferation and measurements based on the mCherry signal that is present in HRasV12^-^ and HRasV12^+^ brains revealed an increased brain size in HRasV12^+^ larvae from 3 dpf onwards (Mayrhofer et al., 2017). Our own brain volume measurements confirmed this, and we observed a significant increase of the larval brain volume at 5 dpf in HRasV12^+^ brains (5.4 ± 0.8 10^6^ μm^3^) compared to HRasV12^-^ brains (2.7 ± 0.3 10^6^ μm^3^) (P < 0.0001) (Fig1B).

**Figure 1:**
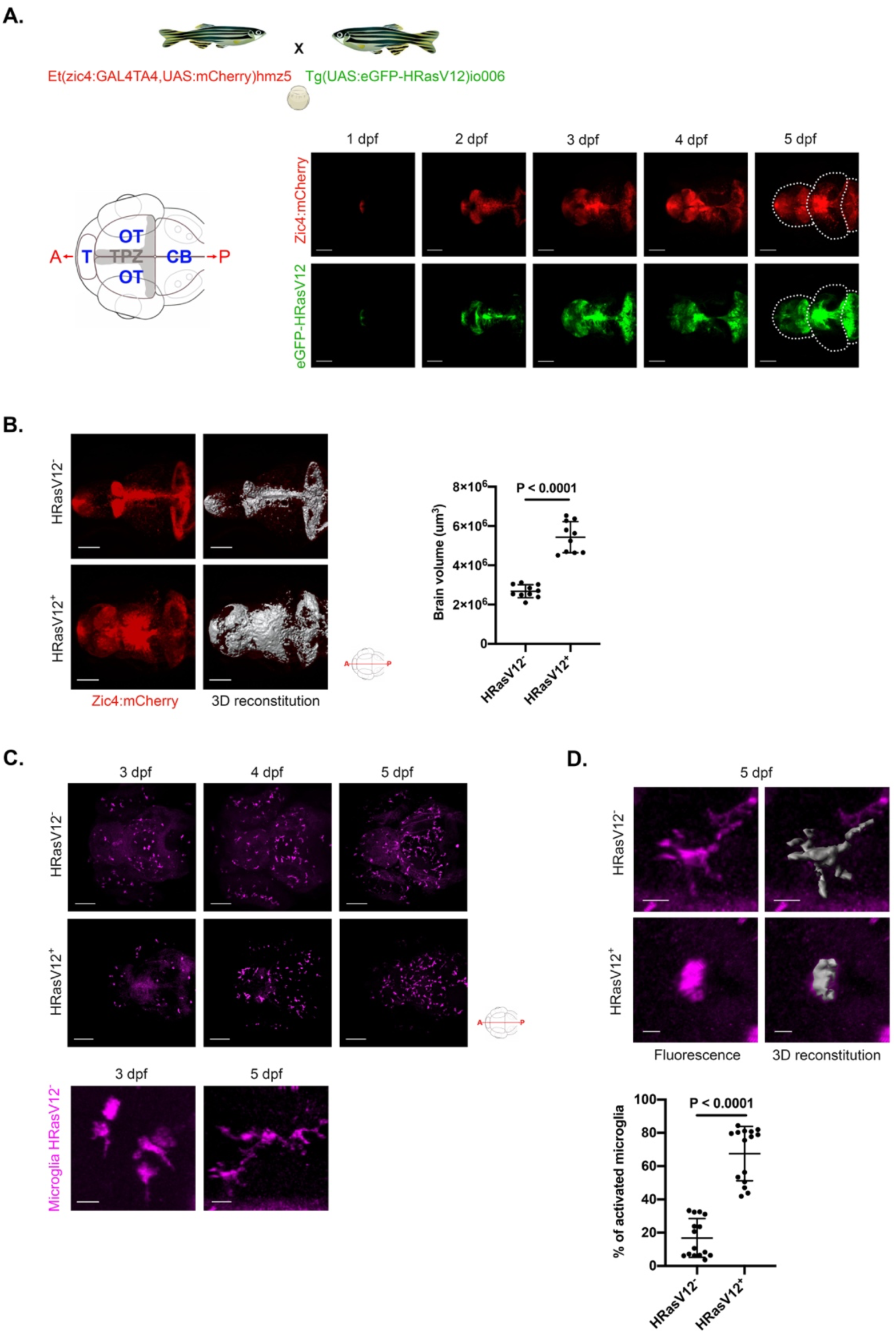
HRasV12 expression in the proliferating regions of the developing CNS alters microglia morphology. (A) Schematic representation of the zebrafish germline system used to induce HRasV12 expression based on the outcross of the indicated fish lines. Schematic anterior-posterior dorsal view of the brain representing the main sub-divisions: telencephalon (T), optic tectum (OT) cerebellum (CB) and tectal proliferation zone (TPZ) in grey. Confocal images showing mCherry and eGFP- HRasV12 fluorescent signal in the proliferating regions of the developing brain of HRasV12^+^ larvae from 1 to 5 dpf. White dotted lines mark the main brain sub-divisions. Scale bar represents 100 μm. (B) Brain volume was assessed using Imaris surface tool to build the segmented images (right panels) of the mCherry signal (left panels) of proliferating regions of the developing brain from 5 dpf HRasV12^-^ (top panels) and HRasV12^+^ (bottom panels). Scale bar represents 100 μm. Brain volumes of 5 dpf HRasV12^-^ and HRasV12^+^ larvae were quantified. HRasV12^-^: n = 10; HRasV12^+^: n = 10; N = 3. (C) Confocal images of microglia (magenta) distribution throughout the developing brain of HRasV12^-^ (top panels) and HRasV12^+^ brains (bottom panels) from 3 to 5 dpf, using 4C4 antibody to specifically label microglia. Scale bar represents 100 μm. Close-ups of microglia at 3 dpf and 5 dpf under physiological condition (HRasV12^-^) allow to determine their morphology: amoeboid (activated) and ramified (surveillance). Scale bar represents 10 μm. (D) Close-ups of microglia from 5 dpf HRasV12^-^ (top panels) and HRasV12^+^ (bottom panels) larvae (left panels) and their segmented images in 3D (right panels) using Imaris surface tool, to assess microglia morphology. Scale bar represents 10 μm. The number of activated microglia was quantified within the microglial population of 5 dpf control and HRasV12^+^ larvae. Results are shown as a percentage of total microglia. HRasV12^-^: n = 15; HRasV12^+^: n = 15; N = 3. Error bars represent mean ± SD. Images were captured using a Zeiss LSM710 confocal microscope with a 20X/NA 0.8 objective. All images represent the maximum intensity projections of Z stacks.

To visualise microglia, we performed immunohistochemistry using the microglia specific zebrafish 4C4 antibody in HRasV12^-^ (control) and HRasV12^+^ larvae (Becker and Becker, 2001; Chia et al., 2018; Mazzolini et al., 2019; Tsarouchas et al., 2018). In line with normal zebrafish development, we did not detect microglia progenitor cells in the brain at 1 and 2 dpf (data not shown), whereas they appear within the tectum by 3 dpf, then spread to the entire brain during development in both conditions (Fig1C). Under normal physiological conditions, mature microglia are highly ramified and are constantly scanning their microenvironment (Nimmerjahn et al., 2005). Once microglia detect changes, such as microenvironment modifications in brain pathologies, they retract their processes and adopt an amoeboid morphology which reflects their activation (Karperien, 2013). During zebrafish development microglia show an amoeboid phenotype during brain colonisation stages at 3 dpf and transit towards a ramified phenotype over the next two days of development (Mazzolini et al., 2019; Svahn et al., 2013) (Fig1C). We previously reported an activated microglial morphology in the presence of pre-neoplastic AKT1 neural cells in larval zebrafish brains (Chia et al., 2018). In order to test if the activated morphology is a general feature upon exposure to pre-neoplastic cells, we analysed microglia morphology in HRasV12^+^ larvae at 3 dpf and upon exposure to HRasV12^+^ cells for 2 days (5 dpf). To assess microglial morphology, we used our previously described method to calculate the ratio of the microglia cell surface to microglia cell volume, which can be used as a read out for their activation (Chia et al., 2018) (Fig1D). Within 3 dpf HRasV12^-^ and HRasV12^+^ brains, microglia were mostly amoeboid in both conditions with a percentage of activation of 80% ± 4 and 84% ± 5 respectively (Sup1A). However, in 5 dpf larvae we detected 67 ± 16 % of activated microglia in presence of HRasV12^+^ cells compared to 17 ± 12 % in the HRasV12^-^ condition (P < 0.0001) (Fig1D). These results show that the presence of pre-neoplastic cells, independent of oncogene and cell type, affects the establishment of microglia ramification and confers them with an active phenotype.

Microglia numbers have been shown to be increased in late stage tumours but also during tumour initiation (Badie and Schartner, 2001; Bowman et al., 2016; Chen et al., 2012; Chia et al., 2018; Coniglio and Segall, 2013; Graeber et al., 2002; Li and Graeber, 2012). To test whether the expression of HRasV12 in the developing CNS induces an increase in microglial numbers, we quantified 4C4^+^ cells (microglia) within 3 and 5 dpf brains of control and HRasV12^+^ larvae. Interestingly, while microglial numbers were similar in both conditions at 3 dpf,103 ± 19 and 93 ± 19 microglia respectively (Sup1B), we detected a significant increase in microglial numbers in HRasV12^+^ brains at 5 dpf with 198 ± 34 microglia compared to 129 ± 20 microglia in control brains (P < 0.0001) (Fig2A).

**Figure 2:**
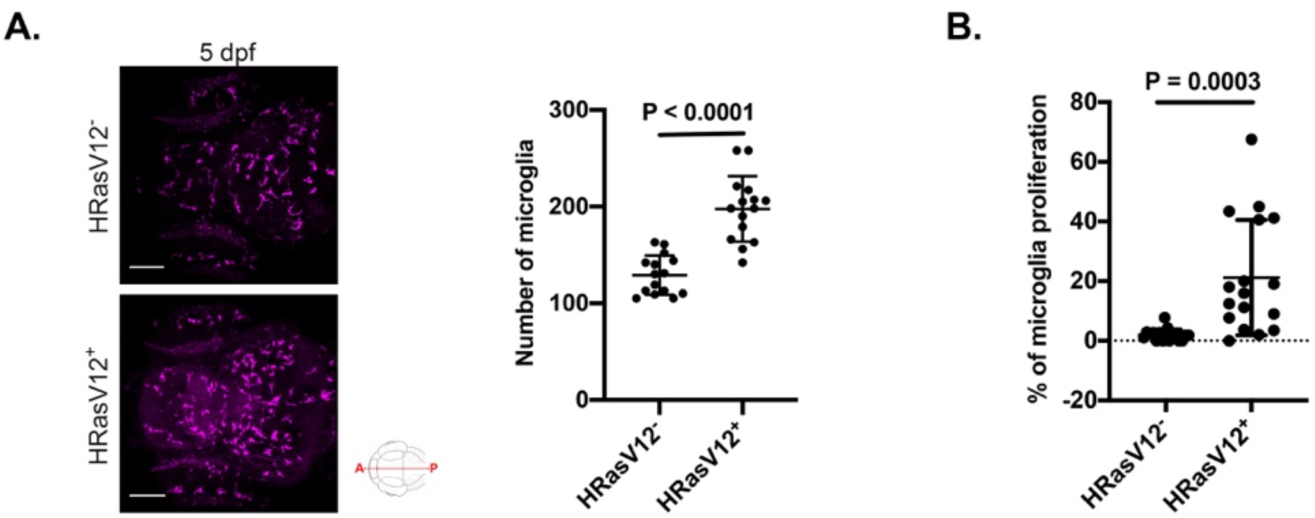
Pre-neoplastic cells promote microglia proliferation. (A) Confocal images of the microglial population (magenta) of 5 dpf HRasV12^-^ (top panel) and HRasV12^+^ (lower panel) larvae. Scale bar represents 100 μm. Quantifications revealed a higher number of microglia in HRasV12^+^ brains compared to HRasV12^-^ brains at 5 dpf. HRasV12^-^: n = 15; HRasV12^+^: n = 15; N = 3. Error bars represent mean ± SD. (B) To measure microglia proliferation, the number of 4C4^+^/EdU^+^ cells was measured within the microglial population of 5 dpf control and HRasV12^+^ larvae. Results are expressed as a percentage of total microglia. HRasV12^-^: n = 17; HRasV12^+^: n = 17; N = 3. Error bars represent mean ± SD. Error bars represent mean ± SD. Images were captured using a Zeiss LSM710 confocal microscope with a 20X/NA 0.8 objective. All images represent the maximum intensity projections of Z stacks.

To understand how this increase of the microglia population is achieved, we assessed microglial proliferation activity by performing an EdU assay, and stained control and HRasV12^+^ larval microglia (4C4^+^). The number of EdU^+^/4C4^+^ cells was significantly higher in HRasV12^+^ brains (21 ± 19 %) compared to control brains (2 ± 2 %) (P = 0.0003) (Fig2B). Thus, the exposure of microglia to pre-neoplastic HRasV12^+^ cells from 3 to 5 dpf triggers their proliferation and results in a significant increase of their number.

Taken together, our results show that microglia number and morphology are not affected by the immediate presence of pre-neoplastic cells, however within 2 days, microglia numbers increased significantly, and showed a higher percentage of activation compared to controls.

### Pre-neoplastic HRasV12^+^ cells affect microglial functions

In light of our previous observations on activated microglial morphology, we decided to investigate effects of HRasV12^+^ cells on fundamental microglial functions such as phagocytosis and motility. The clearance of dying cells and debris in the brain is carried out by microglia and their aptitude to respond to lesions is well described in the literature (Ayata et al., 2018; Davalos et al., 2005; Neumann et al., 2008; Nimmerjahn et al., 2005; Norris et al., 2018; Peri and Nüsslein-Volhard, 2008; Sieger et al., 2012). To measure effects on microglial phagocytic activity, we developed a phagocytosis assay based on zymosan injection (Fig3A). Zymosan is a yeast cell wall well described to trigger macrophage phagocytosis involving CR3, Mannose, Dectin- 1 and Toll-like receptors (Bruggen et al., 2009; Cabec et al., 2000; Mazzolini et al., 2010; Underhill and Ozinsky, 2002). We injected zymosan coupled with a fluorochrome into either the telencephalon or tectum of 5 dpf control and HRasV12^+^ larval brains, whereas 3 dpf larval brains were only injected into the tectum as very few microglia are detected in the telencephalon at that developmental stage (Fig1C). Upon zymosan injection, larvae were incubated for 6 hours, followed by fixation and immunohistochemistry for microglia using the 4C4 antibody (Fig3A). This assay allowed us to quantify the percentage of phagocytosed zymosan by microglia in HRasV12^-^ and HRasV12^+^ brains to determine whether pre-neoplastic cells affect their phagocytic activity (Fig3A). The percentage of microglial phagocytosis was not significantly different in the tectum of 3 dpf control (19 ± 22 %) and HRasV12^+^ larvae (14 ± 12 %), whereas microglial phagocytosis was significantly reduced in 5 dpf HRasV12^+^ larvae (tectum: 16 ± 20 %; telencephalon: 15 ± 19 %) compared to control larvae (tectum: 31 ± 21 %; telencephalon: 41 ± 22 %) (P_tectum_ = 0.0074) (P_telencephalon_ = 0.0023) (Fig3B). These results indicate that within 2 days of contact with pre- neoplastic cells, microglia exhibit a strong reduction of their phagocytic activity.

**Figure 3:**
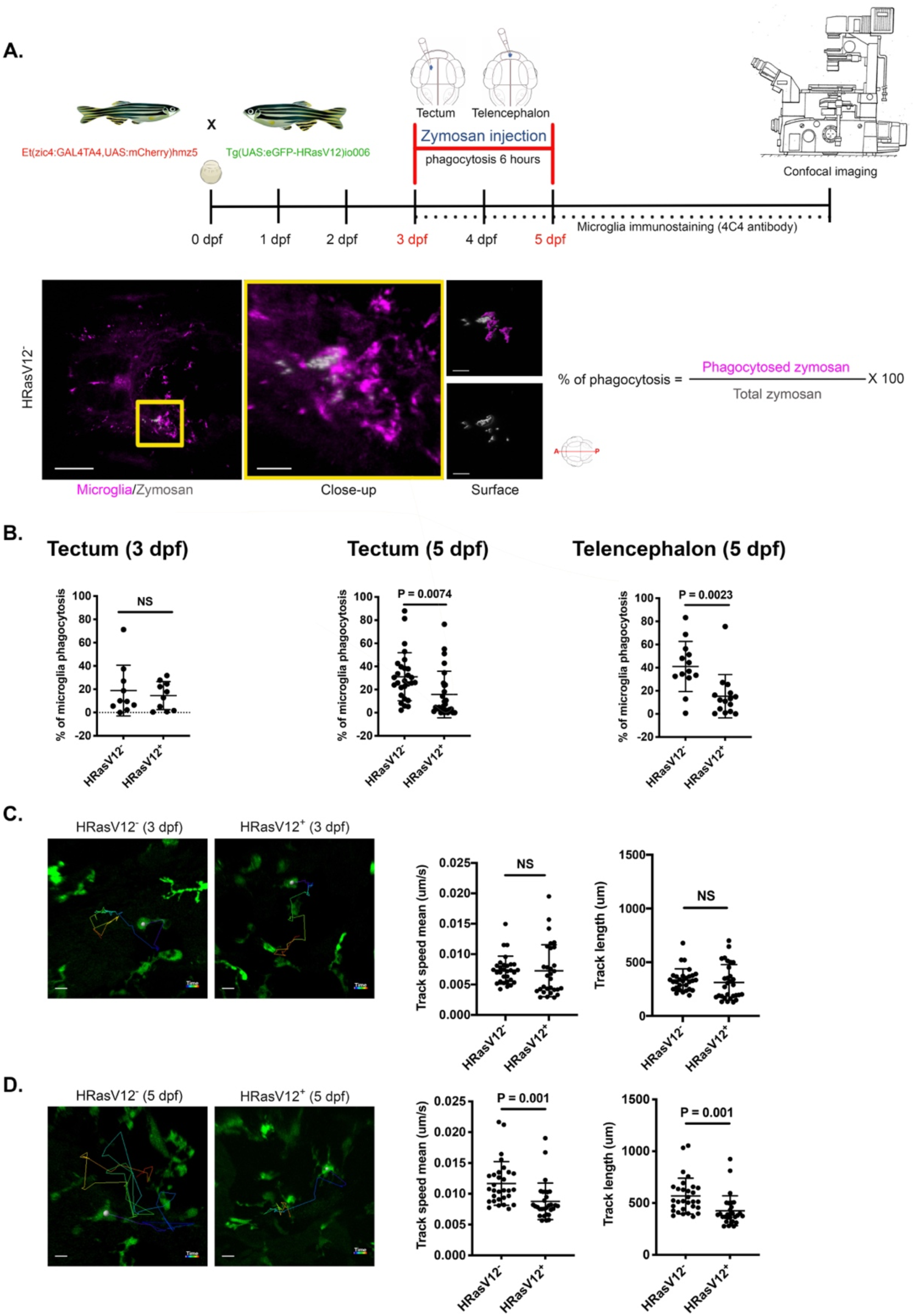
HRasV12^+^ cells affect actin cytoskeleton dependent microglial functions. (A) Schematic representation of the phagocytosis assay used to measure microglia phagocytic activity of 3 and 5 dpf HRasV12^-^ and HRasV12^+^ larvae. Zymosan coupled with a fluorochrome was injected into either the telencephalon or the tectum of HRasV12^-^ and HRasV12^+^ larvae at 3 and 5 dpf. Larvae were incubated for 6 hours post-injection at 28.5 °C, fixed, then labelled with the 4C4 antibody to visualise microglia. Confocal image of a 5 dpf control larval brain injected with zymosan (white) into tectum (yellow square). Scale bar represents 100 μm. Close-up of the injection site reveals zymosan phagocytosed by microglia (magenta). Scale bar represents 20 μm. The Imaris surface tool was used to segment and read out the sum of fluorescence from zymosan internalized by microglia (magenta surface) as well as the total amount of injected zymosan within the telencephalon or tectum (grey surface). The percentage of phagocytosis was calculated following the indicated formula. (B) Efficiency of phagocytosis was calculated for 3 and 5 dpf HRasV12^-^ and HRasV12^+^ larvae injected with zymosan into either the telencephalon or the tectum. Results are expressed as a percentage of the total amount of injected zymosan. 3 dpf: HRasV12^-^ : n = 10; HRasV12^+^: n = 10; N = 3.; 5 dpf (tectum): HRasV12^-^: n = 28; HRasV12^+^: n = 28; N = 3; 5 dpf (telencephalon): HRasV12^-^: n = 13; HRasV12^+^: n = 13; N = 3; Error bars represent mean ± SD. (C-D) Microglia movement in 3D (motility) was tracked using Imaris tracking tool for the full duration of time series (13 hours, *Δ*t = 14 min). Track speed mean and track length were calculated in HRasV12^-^ and HRasV12^+^ conditions at 3 (C) and 5 dpf (D). 3 dpf: HRasV12^-^: n = 30; HRasV12^+^: n = 30; N = 3.; 5 dpf: HRasV12^-^: n = 30; HRasV12^+^ n = 3; N = 3; Error bars represent mean ± SD. Examples of microglia tracks displayed as time color-coded lines from the different conditions. Scale bar represents 20 μm. Images were captured using a Zeiss LSM880 confocal microscope with a 20X/NA 1.0 objective. All images represent the maximum intensity projections of Z stacks.

During early stages of brain development, microglia are very efficient at clearing debris and apoptotic cells due to their high motility (Haynes et al., 2006; Kyrargyri et al., 2020; Madry et al., 2018b; Sieger et al., 2012). To assess if the observed change in microglial morphology/function in HRasV12^+^ larvae has an impact on their motility, we performed high resolution confocal live imaging on HRasV12^+^ larvae from the outcross between *Tg(zic4:Gal4UAS:mCherry:mpeg1:eGFP)* and *Tg(UAS:TagBFP2-HRasV12)* fish lines, and control larvae from the incross between *Tg(zic4:Gal4UAS:mCherry:mpeg1:eGFP)*. The expression of eGFP under the mpeg1 promoter allows to visualise all macrophages including microglia (Ellett et al., 2011). Based on our previous observations on microglia morphology, number and phagocytosis, we decided to perform confocal live imaging for 13 hours on larvae between 3 and 4 dpf and between 4 and 5 dpf. We tracked microglial movement in 3 dimensions (3D) for the full duration of the time series and calculated the track length (motility) and their speed of movement in the two different conditions. Interestingly, in presence of HRasV12^+^ cells at 3 dpf, microglia speed (0.007 ± 0.002 μm/s) and motility (312 ± 165 μm) were similar to control microglia speed (0.007 ± 0.004 μm/s) and motility (335 ± 103 μm) (Fig3C). However, we observed an obvious reduction of microglia motility in presence of pre-neoplastic cells at 5 dpf compared to control (Fig3D). Quantification of microglial speed (0.009 ± 0.003 μm/s) and motility (425 ± 144 μm) in HRasV12^+^ larvae compared to speed (0.012 ± 0.003 μm/s) and motility of microglia in HRasV12^-^ larvae (567 ± 173 μm) showed a significant difference (P_speed_ = 0.001) (P_motility_ = 0.001) (Fig3D).

In summary, our results show that within a short time window, pre-neoplastic cells alter not only microglial morphology but also key functions such as phagocytosis and motility at early stages of GBM formation. Hence, we speculated that lasting alterations of the microglial actin cytoskeleton might be the underlying cause, which result in a permanent change in morphology and impair related functions such as motility and phagocytosis.

### RNA sequencing of isolated microglia reveals down-regulation of *wasla* gene expression

Our data show that microglia morphology, phagocytosis and motility are altered upon contact with pre-neoplastic cells. These cellular mechanisms are regulated by a compilation of different signalling pathways, but they are all orchestrated by the actin cytoskeleton organisation (Freeman and Grinstein, 2014; Pollard and Cooper, 2009; Svitkina, 2018). Hence, we decided to conduct RNA sequencing to investigate differentially expressed (DE) genes involved in actin cytoskeleton organisation and regulation between control and HRasV12^+^ conditions at 3 and 5 dpf. Following our previously published protocols we isolated microglia from dissociated brains and performed 4C4 immunohistochemistry followed by fluorescence-activated cell sorting (FACS) (Mazzolini et al., 2018, 2019) (Sup2). For each time point, microglia were pooled from 600 HRasV12^-^ and HRasV12^+^ larval brains and three replicates were performed per time point for RNA sequencing.

We evaluated the expression correlation between biological replicates using the whole set of genes from the RNA-seq data set. Each sample consisted of isolated microglia from 600 brains; of note, scatter plots of the normalized transformed read counts showed that biological replicates were highly correlated (r > 0.75). Interestingly, correlation between control samples was higher at 3 dpf (r > 0.83) and 5 dpf (r > 0.81) than between HRasV12^+^ samples (r > 0.75) and (r > 0.79) (Sup3A). The lower correlation obtained at 3 and 5 dpf from HRasV12^+^ larval brains could be explained by heterogeneity in those samples due to the presence of pre-neoplastic cells (Sup3A). Principal component analysis (PCA) confirmed this correlation by showing HRasV12^-^ replicates more clustered than replicates from HRasV12^+^ conditions. Moreover, clusters corresponding to 3 and 5 dpf samples from both conditions were well segregated (Sup3B).

Using the KEGG database, we defined pathways related to actin cytoskeleton in zebrafish (Regulation of actin cytoskeleton, Focal adhesion, Adherens junction, Tight junction and mTOR signalling pathway) and selected genes from our RNA-seq which are referred to them (Table 1). We compared expression of these genes in microglia between control and HRasV12^+^ larval brains at 3 and 5 dpf and focussed on genes that are differentially expressed (FDR < 0.05, fold change > |2|) at 5 dpf when pre- neoplastic cells affect microglial functions. We obtained 11 differentially expressed (DE) genes, 8 with lower expression and 3 with higher expression in microglia from HRasV12^+^ brains compared to control microglia (Fig4A, Table1). The majority of these genes belonged to the “Regulation of actin cytoskeleton” pathway, and the identified top ranked DE gene was *wasla* (Table 1), the zebrafish orthologue of human WASP like actin nucleation promotion factor (WASL, also known as N-WASP). WASL is a key protein of the actin cytoskeleton organisation, necessary to maintain cell shape, efficient phagocytosis and motility (Dogterom and Koenderink, 2019; May et al., 2000; Niedergang and Grinstein, 2018; Yamaguchi et al., 2005). *Wasla* showed 4.2 times lower expression in microglia in the presence of pre-neoplastic cells compared to microglia from control brains (FDR = 0,005). Intriguingly, *wasla* was the only gene of the WASP family significantly differentially expressed in microglia in the presence of pre-neoplastic cells, whereas *was*, *wasf* and *wash* expressions were the same as in control microglia (Fig4B).

**Figure 4:**
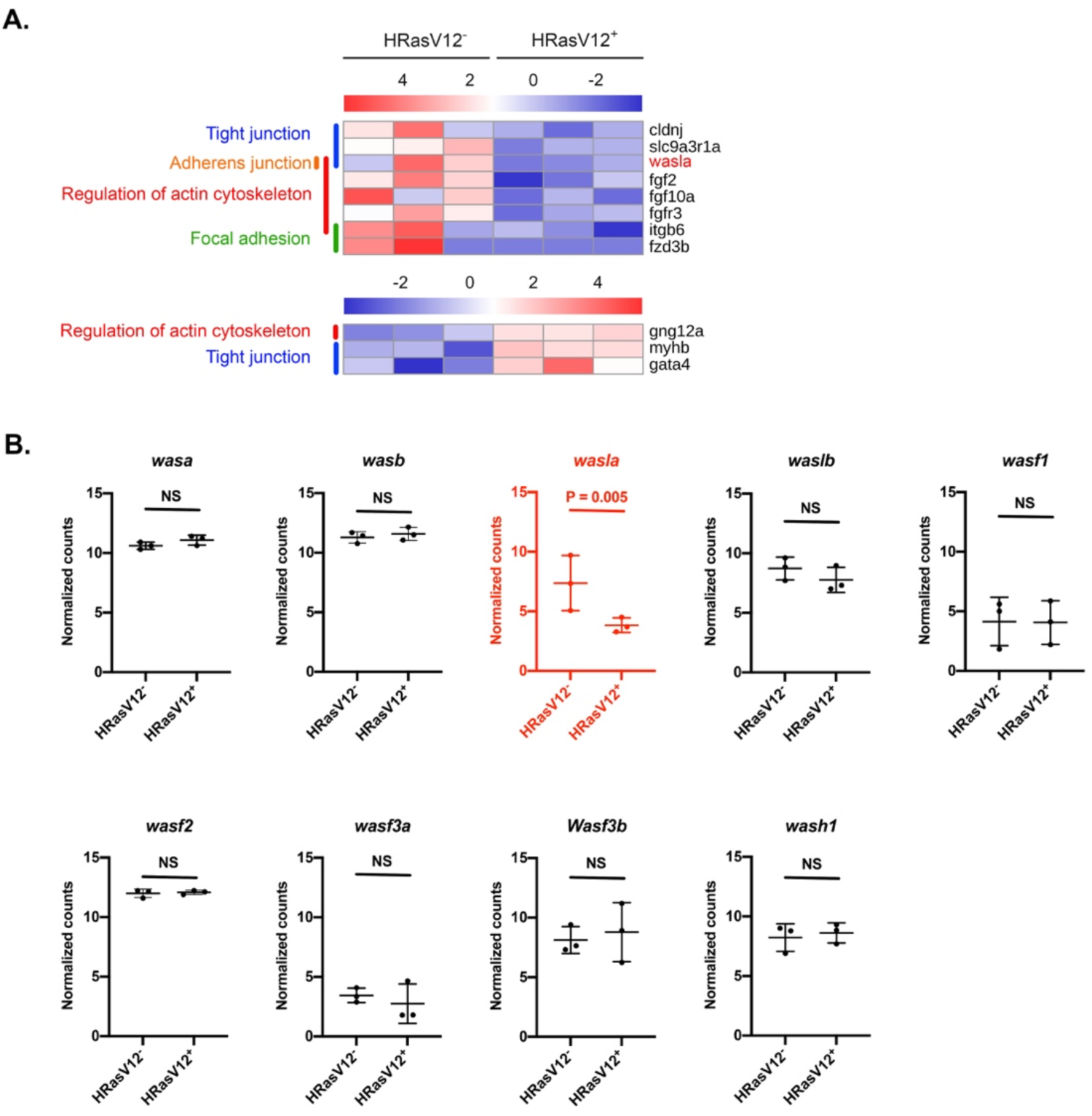
The actin nucleation promoting factor *wasla* is less expressed in microglia from HRasV12^+^ larvae. (A) Heatmap of differentially expressed (DE) genes (FDR < 0.05, Fold Change > |2|) from microglia transcriptome comparisons between 5 dpf HRasV12^-^ and HRasV12^+^ larval brains (11 genes), belonging to zebrafish actin cytoskeleton KEGG pathways [Regulation of actin cytoskeleton, Focal adhesion, Adherens junction, Tight junction and mTOR signalling pathway]. The actin nucleation promoting factor *wasla* is the top ranked gene of the list. See also Table S1. (B) Dot plots of normalized transformed read counts of the representative WASP family genes. Black plots represent non DE genes whereas, red plots correspond to DE genes at 5 dpf between control and HRasV12^+^ conditions. The means ± SD of three independent experiments are plotted. The means ± SD of three independent experiments are plotted.

These results show that microglia exposed to pre-neoplastic cells express a lower level of *wasla*. Thus, we hypothesised that reduced levels of *wasla* are the underlying cause of the change of microglial morphology and the decrease of their motility and phagocytotic capacity.

### Microglia specific overexpression of *wasla* restores microglial functions in HRasV12^+^ brains

In order to test our hypothesis that lower expression levels of *wasla* were responsible for the observed changes in microglial morphology and functions, we performed cell specific overexpression for *wasla* in microglia. To achieve this, we created a plasmid which encodes for *wasla* under the control of the mpeg1 promoter specific to microglia and macrophages (Ellett et al., 2011). Injection of the mpeg1:wasla plasmid into one cell stage embryos resulted in a transient, mosaic expression of *wasla* in microglia. To verify the efficiency of this strategy, we injected the plasmid into one cell stage *Tg(mpeg1:mCherry*) embryos, isolated mCherry^+^ microglia/macrophages at 5 dpf by FACS (Sup4A) and performed qPCR for *wasla*. We obtained a 1.46 times higher expression of *wasla* in mCherry^+^ microglia/macrophages from injected embryos compared to non-injected embryos (Sup4B). Hence, we injected the mpeg1:wasla plasmid into HRasV12^-^ and HRasV12^+^ embryos and analysed the impact on microglia at 5 dpf. Injection into HRasV12^-^ embryos (hereafter HRasV12^-^/wasla), did not alter microglia morphology (HRasV12^-^/wasla: 9 ± 4 %; HRasV12^-^ : 7 ± 2 %) and number (HRasV12^-^/wasla: 146 ± 17; HRasV12^-^ : 143 ± 16) compared to control larvae (Sup4C). This shows that higher expression levels of *wasla* do not change microglial appearance under physiological conditions. However, injection of the mpeg1:wasla plasmid into HRasV12^+^ embryos had a significant impact and restored microglia ramification, number, phagocytic activity and motility in this condition. Upon *wasla* overexpression in microglia of HRasV12^+^ larvae, their activation was significantly reduced in HRasV12^+^/wasla (33 ± 14 %) compared to the HRasV12^+^ condition (67 ± 16 %) (P < 0.0001) and their morphology similar to control microglia (Fig5A, top panels; Fig 5B). Furthermore, microglia numbers were significantly reduced compared to the HRasV12^+^ condition (198 ± 34) (P < 0.0001) and were similar to control microglia numbers with 128 ± 32 microglia in HRasV12^+^/wasla compared to 131 ± 22 in control larvae (Fig5A, middle panels; Fig 5C). Finally, microglial phagocytic activity (53 ± 32 %), speed (0.012 ± 0.003 μm/s) and motility (584 ± 156 μm) in 5 dpf HRasV12^+^/wasla larvae were significantly different to HRasV12^+^ larvae (phagocytosis: 16 ± 20 %) (speed: 0.009 ± 0.003 μm/s) (motility: 426 ± 144 μm), and reached same values as control larvae (phagocytosis: 31 ± 21 %) (speed: 0.012 ± 0.003 μm/s) (motility: 567 ± 173 μm) (P_phagocytosis_ < 0.0001) (P_speed_ = 0.0005) (P_motility_ = 0.0006) (Fig5A, lower panels; Fig 5D).

**Figure 5:**
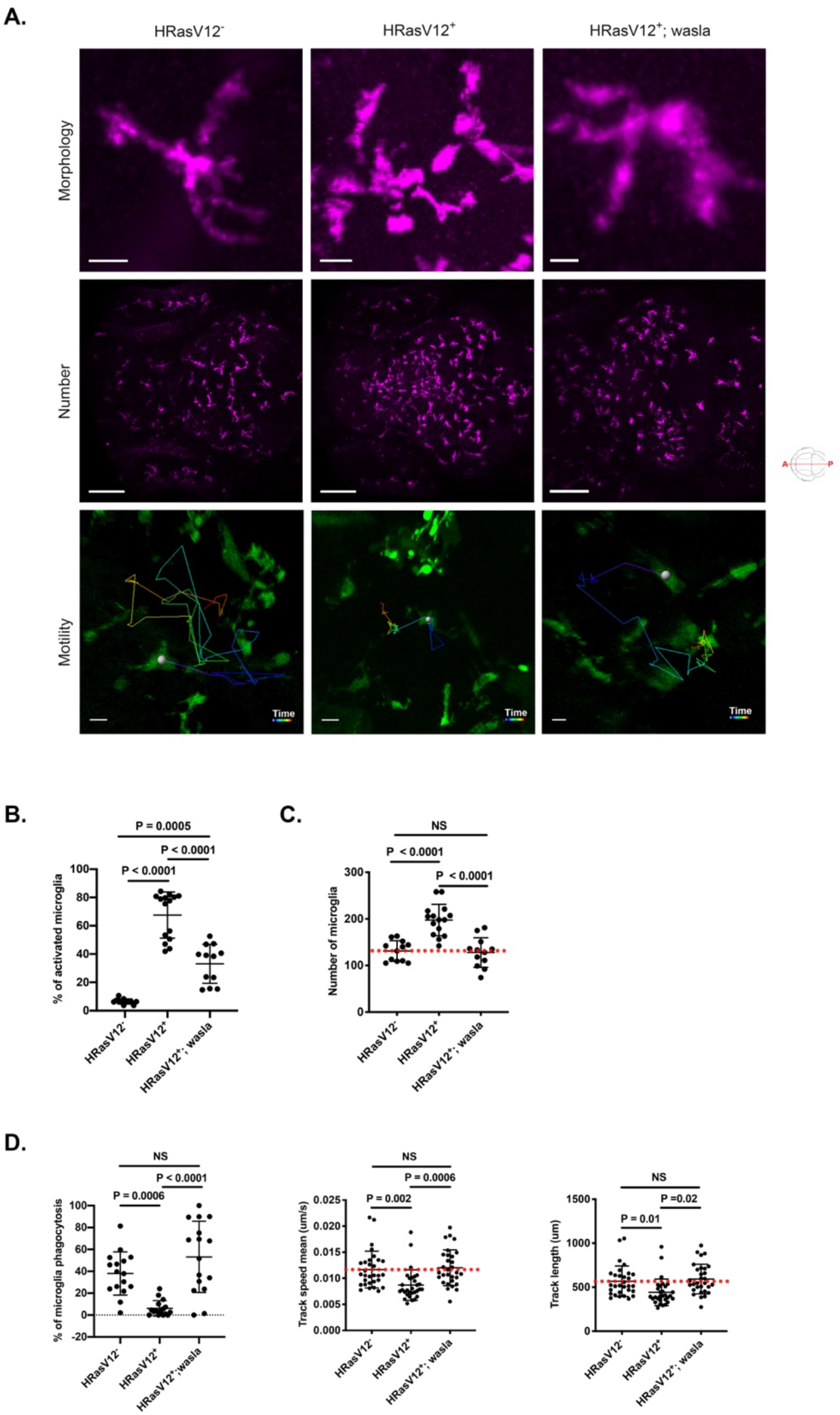
***Walsa* overexpression in microglia restores their morphology and functions.** (A) Close-up confocal images of microglia (top panels), microglial population (middle panels) and examples of microglia tracks displayed as time color- coded lines (lower panels) from 5 dpf HRasV12^-^ (left panel), HRasV12^+^ (middle panel) and HRasV12^+^; wasla (right panel) brains. Scale bar represents 10, 100 and 20 μm respectively. (B) The percentage of activated microglia and (C) the total number of microglia from 5 dpf HRasV12^-^, HRasV12^+^ and HRasV12^+^; wasla brains were quantified. HRasV12^-^: n = 15; HRasV12^+^: n = 15; HRasV12^+^; wasla: n = 15; N = 3. Error bars represent mean ± SD. (D) The percentage of microglia phagocytosis within tectum of 5 dpf HRasV12^-^, HRasV12^+^ and HRasV12^+^;wasla as well as microglia track speed mean and track length (motility) were quantified. Phagocytosis: HRasV12^-^: n = 16; HRasV12^+^: n = 16; HRasV12^+^; wasla: n = 16; N = 3. Motility: HRasV12^-^: n = 30; HRasV12^+^: n = 30; HRasV12^+^; wasla: n = 30; N = 3. Error bars represent mean ± SD. Red dotted lines indicate the number of microglia, track speed mean or track length in control condition. Images were captured using a Zeiss LSM880 confocal microscope with a 20X/NA 0.8 objective. All images represent the maximum intensity projections of Z stacks.

In summary, these results reveal that *wasla* is a key gene to maintain microglial functions and its lower expression levels in HRasV12^+^ brains are responsible for alterations in microglia morphology, phagocytosis and motility.

### Rescue of *wasla* expression in microglia slows down pre-neoplastic growth by restoring an efficient microglial phagocytic activity

Intrigued by the restoration of microglia morphology and functions by *wasla* overexpression in the HRasV12^+^ condition, we investigated its impact on pre- neoplastic growth and survival of the larval zebrafish. The Zic/HRasV12 model has been previously described with a survival rate of 4% in the first month (Mayrhofer et al., 2017). Therefore, we monitored survival in HRasV12^-^, HRasV12^+^ and HRasV12^+^/wasla conditions daily for one month. The survival of HRasV12^+^ larvae reached 7 ± 6 % at 1 month post fertilisation (mpf) compared to 83 ± 16 % for controls, which is in line with previously published data (Fig6A) (Mayrhofer et al., 2017). The number of HRasV12^+^ surviving larvae dropped drastically around 10 dpf, however, HRasV12^+^/wasla larvae showed better survival during these time points and deteriorated with a significant delay (P = 0.0017; Gehan-Breslow-Wilcoxon test). Finally, HRasV12^+^/wasla larvae reached a similar survival rate as HRasV12^+^ larvae (5 ± 5 %) at 31 dpf (Fig6A).

These results show that *wasla* overexpression in microglia improves survival during the initial stages of pre-neoplastic growth in HRasV12^+^ larvae. Hence, we speculated that tumour formation has been altered in HRasV12^+^/wasla larvae. To investigate whether *wasla* overexpression in microglia affects pre-neoplastic mass growth (HRasV12^+^ cells), we measured the brain volume (mCherry signal) and pre-neoplastic mass (eGFP signal) of 5 dpf HRasV12^+^/wasla brains compared to HRasV12^+^ and control brains. Interestingly, *wasla* overexpression in microglia significantly reduced the larval brain volume (1.7 ± 0.4 10^6^ μm^3^; P= 0.055) and pre-neoplastic mass (2.7 ± 0.5 10^6^ μm^3^; P= 0.012) compared to HRasV12^+^ brains (2.3 ± 0.5 10^6^ μm^3^ and 3.6 ± 0.6 10^6^ μm^3^ respectively). Notably, the HRasV12^+^/wasla brain volume was reduced to the same volume as that of control brains (1.7 ± 0.3 10^6^ μm^3^) (Fig6B). These data reveal that the survival improvement of HRasV12^+^/wasla larvae correlates with a significant reduction of their brain volume and pre-neoplastic mass. As we have shown that microglial morphology and actin-dependent functions such as phagocytosis were restored in these larvae, we hypothesised that phagocytosis of pre-neoplastic cells by microglia contributed to the reduced pre-neoplastic growth and better survival. To address this hypothesis, we tested the direct impact of rescued *wasla* expression in microglia on phagosome formation by quantifying the number of phagosomes per microglia of 5 dpf HRasV12^+^ and HRasV12^+^/wasla larval brains. If *wasla* was, as we predicted, involved in microglia actin cytoskeleton organisation then its overexpression should modify phagosome formation (Marion et al., 2012; May et al., 2000; Niedergang and Grinstein, 2018). To quantify the number of phagosomes we performed high resolution confocal imaging on 5 dpf HRasV12^+^/wasla and HRasV12^+^ larvae from the outcross between *Tg(zic4:Gal4UAS:mCherry:mpeg1:eGFP)* and *Tg(UAS:TagBFP2- HRasV12)* fish lines. This model allowed us to easily visualise phagosomes as black holes within microglia as the GFP signal is restricted to the cytoplasm (Fig 6C). Of note, microglia overexpressing *wasla* in HRasV12^+^ larvae contained 3 ± 3 phagosomes whereas HRasV12^+^ microglia contained only 1 ± 1 phagosome (P < 0.0<u>0</u>01) (Fig 6C). Therefore, these results confirm that rescue of *wasla* expression in microglia of HRasV12^+^ larvae increases phagosome formation. Hence, we tested if this leads to an increase in phagocytosis of HRasV12^+^ cells. To investigate the phagocytosis of HRasV12^+^ cells (GFP signal) by microglia, we performed flow cytometry and quantified the mean of fluorescence from GFP^+^ (HRasV12^+^) cells phagocytosed by isolated microglia from HRasV12^+^ and HRasV12^+^/wasla larval brains (Table2). These quantifications revealed that the amount of pre-neoplastic cells engulfed by HRasV12^+^/wasla microglia was 1.5 higher than in the HRasV12^+^ condition (P = 0.0002) (Fig6D). Hence, the restoration of phagocytic activity in microglia results in an increased phagocytosis of pre-neoplastic HRasV12^+^ cells. This might explain the smaller pre-neoplastic mass volume in HRasV12^+^/wasla larvae.

**Figure 6:**
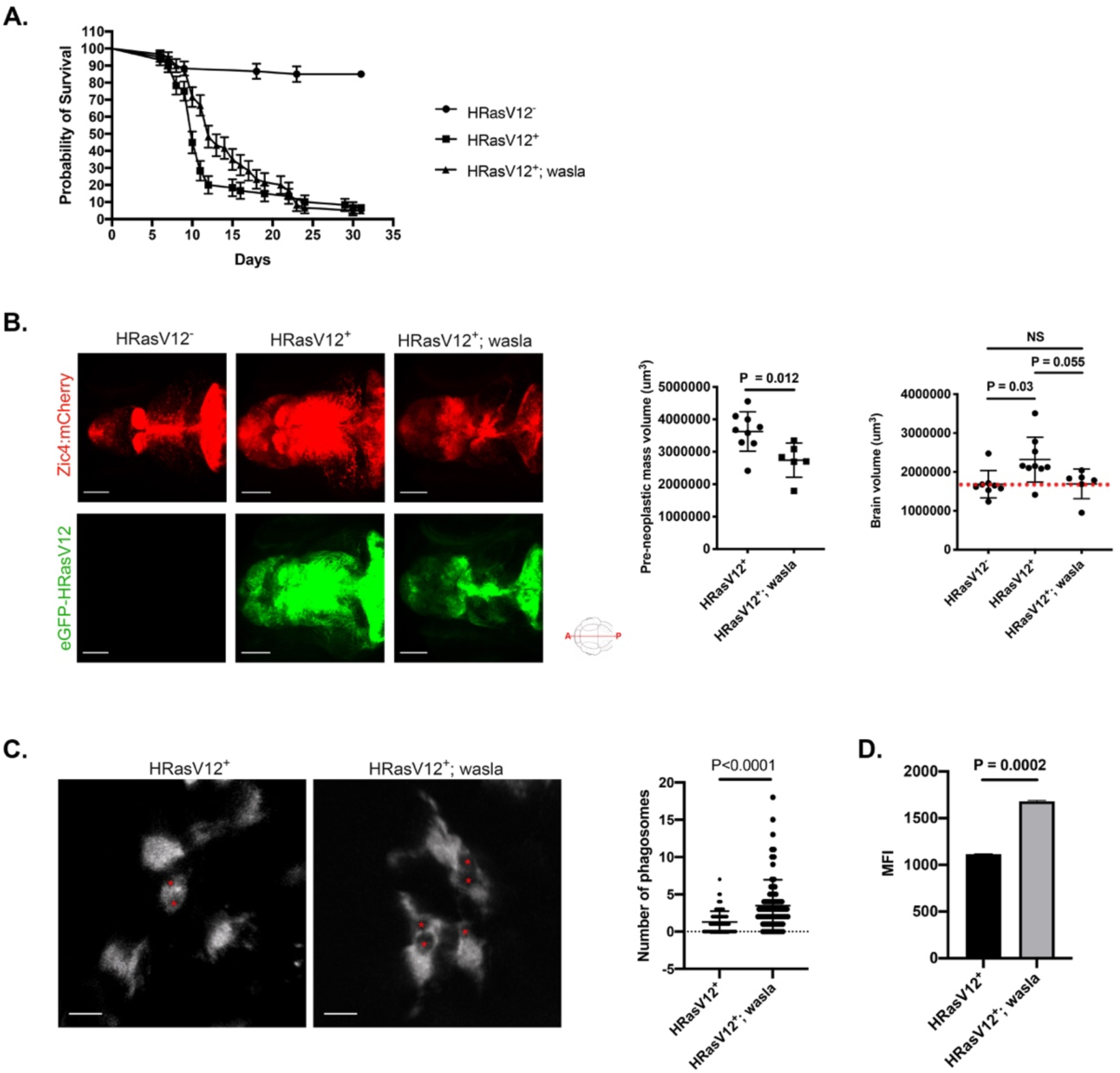
Microglial *wasla* expression is crucial to slow down tumour progression. (A) Kaplan-Meier survival plot of HRasV12^-^, HRasV12^+^ and HRasV12^+^;wasla larvae control over 31 days, n = 50/60, 4/60 and 3/60 respectively. P = 0.0017 (Gehan-Breslow-Wilcoxon test between HRasV12^+^ and HRasV12^+^;wasla conditions). Error bars represent mean ± SD. (B) Brain and pre-neoplastic mass volume were measured using the mCherry signal (brain, top panels) and eGFP signal of HRas^+^ cells (pre-neoplastic mass; bottom panels) of proliferating regions of the developing brain from 5 dpf HRasV12^-^ (left panel), HRasV12^+^ (middle panel) and HRasV12^+^;wasla (right panel) larvae. Scale bar represents 100 μm. Brain and pre- neoplastic mass volume from 5 dpf HRasV12^-^, HRasV12^+^ and HRasV12^+^;wasla larvae are quantified using Imaris surface tool. HRasV12^+^: n = 9; HRasV12^+^; wasla: n = 6; N = 3. Error bars represent mean ± SD. Red dotted line indicates the brain volume mean in control condition. (C) Close-up confocal images of microglia (mpeg1:eGFP^+^ cells) from 5 dpf HRasV12^+^ (left panel) and HRasV12^+^; wasla (right panel) brains. Phagosomes are indicated by red asterisks. Scale bar represents 10 μm. The number of phagosomes per microglia from 5 dpf HRasV12^+^ and HRasV12^+^; wasla brains were quantified. HRasV12^+^: n = 80; HRasV12^+^; wasla: n = 80; N = 3. Error bars represent mean ± SD. (D) Mean of GFP fluorescent intensity from phagocytosed pre-neoplastic cells detected by flow cytometry within isolated microglia from 5 dpf HRasV12^+^ and HRasV12^+^;wasla larvae. The means ± SD of two independent experiments are plotted. Images were captured using a Zeiss LSM880 confocal microscope with a 20X/NA 0.8 objective. All images represent the maximum intensity projections of Z stacks.

## Discussion

In this study, we revealed the impact of pre-neoplastic cells on the microglial population, their morphology and functions during tumour initiating stages. Several elegant studies have shown crosstalk between glioma associated microglia (GAMs) and neoplastic cells in the brain creating a microenvironment favourable to tumour growth and maintenance (for review see (Gutmann and Kettenmann, 2019). However, while the activated microglial morphology has been described across models and species (Annovazzi et al., 2018; Bayerl et al., 2016; Chia et al., 2018; Juliano et al., 2018; Kvisten et al., 2019; Resende et al., 2016; Ricard et al., 2016), the underlying causes and the timing for the change in morphology are not understood. To our knowledge, this is the first study to provide evidence of an alteration of microglia morphology and functions due to lower expression levels of the *wasl* gene in presence of tumour initiating cells. We utilised a well-established larval zebrafish brain tumour model to address the earliest stages of tumour induction due to activation of oncogenes. By overexpressing a constitutively active form of the human *HRas* gene, we induced cellular alterations in the larval zebrafish brain that lead to the formation of tumours similar to the mesenchymal subtype of human GBM by one month post fertilization (Mayrhofer et al., 2017). Mesenchymal subtype GBMs have been found to correlate with a stronger enrichment of GAMs compared to proneural and classical GBM subtypes (Bhat et al., 2013; Wang et al., 2017). Here, we strategically worked with 3 and 5 dpf larvae to monitor the pre-neoplastic cell impact on microglia. By using immunohistochemistry, transgenic zebrafish lines for microglia, functional assays and *in vivo* imaging we revealed that increased microglial numbers, morphological changes, reduced motility and phagocytic activity are established at the earliest stages of tumour development. The large number of microglia observed in 5 dpf HRasV12^+^ larval brains resulted from a strong proliferative activity. Abels et al. showed that glioma cells can reprogramme microglia and promote their proliferation by reducing expression levels of *Btg2* gene (Abels et al., 2019). In our transcriptomic data *Btg2* and other genes mediating cell proliferation were not differentially expressed (not shown), an observation that doesn’t correlate with our EdU results confirming increased proliferation. However, the EdU assay provided a readout of all proliferative microglia between 3 and 5 dpf, whereas transcriptomic data have been obtained from a specific timepoint and could reflect only a small population of microglia proliferating at that time as all microglia are not synchronised. A single cell RNA sequencing approach might be suited to circumvent this limitation. Furthermore, as we analysed the microglia population during development a certain degree of proliferation is present even in control larvae, hence the additional increase caused by HRasV12^+^ pre-neoplastic cells might not be strong enough to result in significant changes in gene expression. This could explain why we didn’t detect variation of proliferation gene expression. Within our transcriptomic data, *wasla* was the top ranked DE gene belonging to the “Regulation of actin cytoskeleton” pathway. Interestingly there seems to be controversy on the role of WASL during cell division. Cytokinesis is the final step of cell division taking place at the end of mitosis, this mechanism is characterized by the formation of a contractile actomyosin ring necessary for the separation of the newly forming daughter cells. Wang et al. concluded that WASL has a role in cytokinesis during porcine oocyte maturation (Wang et al., 2020), whereas others consider that mechanism as WASL independent (Bompard et al., 2008; Deschamps et al., 2013; Schwayer et al., 2016). Our data support the hypothesis that WASL is not involved in cell proliferation but is crucial for other microglial functions.

Across species and glioma models, microglia exhibit an “activated” phenotype characterised by an amoeboid shape. The term “activation” encompasses anti- and pro-tumoral microglia and needs more investigation to determine the correlation between microglia polarization and this specific morphology. Nevertheless, this phenotype is a reliable readout of physiological changes within the brain microenvironment. According to Karperien et al. amoeboid microglia are usually characterized by their high capacity to engulf and migrate (Karperien, 2013). However, our results reveal a reduced motility, speed and phagocytic activity of amoeboid microglia upon exposure to pre-neoplastic cells. Our data is in line with a study by Voisin et al. who have co-cultured human microglia cell line (CHME-5) with C6-glioma cells and reported a significant reduction of microglia phagocytic activity after 24 hours exposure (Voisin et al., 2010). In HRasV12^+^ larvae, microglia showed alterations of their functions 2 days after exposure to pre-neoplastic cells. Microglial shape, motility and phagocytic activity are actin-dependent and showed dependency on *wasla*. The rescue of *wasla* expression in microglia of HRasV12^+^ brains restored all these physiological functions and promoted phagocytosis of pre-neoplastic cells. The phagocytosis recovery of microglia correlated with a significant diminution of the pre- neoplastic mass volume; hence we concluded that at 5 dpf microglia efficiently clear pre-neoplastic cells and thus limit their growth. The survival curve supports this observation and showed that overexpression of *wasla* in microglia slowed down the deadly effect of tumour growth on larvae from 5 to 20 dpf. These results are promising but further studies are needed to understand the temporary nature of this effect. One explanation might be the transient and mosaic type of expression generated by the injection of *wasla* constructs into oocytes. Here, not all microglia express sufficient levels of *wasla* and expression diminishes over time. A stable transgenic line expressing high levels of *wasla* in all microglia for a prolonged time would be needed to understand if rescued *wasla* expression can maintain phagocytosis of neoplastic cells by microglia in the long run. However, even under these circumstances, phagocytosis might decrease at later stages of tumour growth due to increased expression of ‘don’t eat me’ signals such as CD47 by tumour cells (Gholamin et al., 2017; Hutter et al., 2019; Li et al., 2017; Ma et al., 2019).

Interestingly, amongst the differentially expressed genes extracted from our transcriptomic data belonging to KEGG pathways linked to “Regulation of actin cytoskeleton” some of the other genes might also contribute to the observed effects. In presence of pre-neoplastic cells microglia expressed lower levels of fibroblast growth factor 2 (fgf2) and fibroblast growth factor receptor 3 (fgfr3), which have been shown to increase microglial migration and phagocytic activity (Noda et al., 2014). Furthermore, *fibroblast growth factor 10a (fgf10a)* showed lower expression in microglia of HRasV12^+^ larvae. FGF10 treatment has been shown to inhibit microglial pro-inflammatory cytokine secretion and proliferation via regulation of the TLR4/NF- _K_B pathway in an animal model after spinal cord injury (Chen et al., 2017). Hence, the reduced expression levels of this gene could contribute to the increased microglial proliferation. Among the DEG that showed higher expression levels in microglia from HRasV12^+^ brains, we detected *G protein subunit gamma 12 (gng12).* Interestingly, *gng12* is known to be highly expressed after LPS stimulation of microglia and to offset the inflammatory response by reducing levels of nitric oxide and TNF*α* (Larson et al., 2010). Moreover, long non-coding gng12 RNAs are highly expressed in glioma tissues and its downregulation inhibits proliferation, migration and epithelial-mesenchymal transition of glioma cells (Xiang et al., 2020). These results suggest that high expression levels of *gng12* in microglia exposed to pre-neoplastic cells might contribute to the generation of a pro-tumoural response.

Phagocytic events are associated with cytokine secretion as part of the innate immune response, and in preparation for adaptive immunity (Acharya et al., 2020; Chung et al., 2006; Fu et al., 2014; Heo et al., 2015; Murray et al., 2005). The secretion of cytokines relies on cellular exocytic pathways involving WASL (González-Jamett et al., 2017; Li et al., 2018; Olivares et al., 2014; Ory and Gasman, 2011). Hence, cytokine secretion of microglia might be altered due to the observed low expression levels of *wasla*. Interestingly, microglia isolated from HRasV12^+^ larval brains showed increased expression levels of *il4* and *il11* (not shown), suggesting the generation of an anti-inflammatory state during tumour initiating stages. Therefore, the assessment of microglial cytokine secretion in HRasV12^+^ and HRasV12^+^/wasla larvae could provide a better understanding of the establishment of an immunosuppressive environment by tumour initiating cells linked to the reduced phagocytic activity of microglia.

To precisely determine the changes in microglial functions in presence of HRasV12^+^ cells, it is important to understand the strategy applied by these cells to alter microglia gene expression levels. Extracellular vesicles (EVs) contain proteins, lipids and different RNA species that change the activity of recipient cells (Tkach and Théry, 2016; Verweij et al., 2019). Several studies have shown the implication of EVs secreted by tumour cells on microglia/macrophages (Abels et al., 2019; Hyenne et al., 2019; Vos et al., 2015). Of note, *wasl* expression has been shown to be regulated by various miRNAs (Bettencourt et al., 2013; Schwickert et al., 2015). Data from the Mione laboratory shows that some of these miRNAs have significantly increased expression levels in HRasV12^+^ cells compared to control cells (Anelli et al., 2018). This includes the miRNA Let-7g-1, which targets the *wasla* gene. Thus, it is tempting to speculate that EVs secreted by HRasV12^+^ cells transfer miRNAs which mediate changes of microglial gene expression and related functions. Future studies will reveal if Let-7g-1 and other miRNAs transferred to microglia via EVs are the underlying cause of the observed changes.

In conclusion, we show for the first time that during tumour initiation stages, pre- neoplastic cells influence microglial functions by altering their gene expression profiles resulting in an alteration of microglial morphology and related functions. We identify *wasla* as a key component in the regulation of microglial morphology, phagocytosis and migration. Our findings provide a mechanism that empowers pre-neoplastic cells to trap microglia within their vicinity, de-activate their phagocytic functions and promote the generation of an anti-inflammatory tumour promoting microenvironment.

## Acknowledgments

This work was supported by a Cancer Research UK Career Establishment Award to Dirk Sieger (C49916/A17494). The authors thank Dr. Marina Mione (Department of Cellular, Computational and Integrative Biology - CIBIO, University of Trento, Trento, Italy) for help and discussion on the project, the BRR zebrafish facility (QMRI, The University of Edinburgh) for maintenance and care of the zebrafish, the SURF Biomolecular core, and the QMRI Flow Cytometry and Cell Sorting Facility. Thanks to SMART-Servier Medical Art for graphic material that has been slightly modified in part (https://creativecommons.org/licenses/by/3.0/). Thanks to Katy Marshall-Phelps for proofreading the manuscript.

## Author contributions

Conceptualisation, J.M. and D.S.; Methodology, J.M. and D.S.; Formal Analysis, J.M., S.LC. and C.C.; Investigation, J.M. and G.M.; Resources, D.S. and JF.Z., Data Curation, J.M., S.LC. and C.C.; Writing – Original Draft, J.M. and D.S., Writing - Review & Editing, J.M., S.LC., JF.Z. and DS; Visualisation, J.M. and S.LC.; Supervision, D.S. and JF.Z., Project Administration, J.M. and D.S.; Funding Acquisition, D.S.

## Supplemental information

**Sup1:**
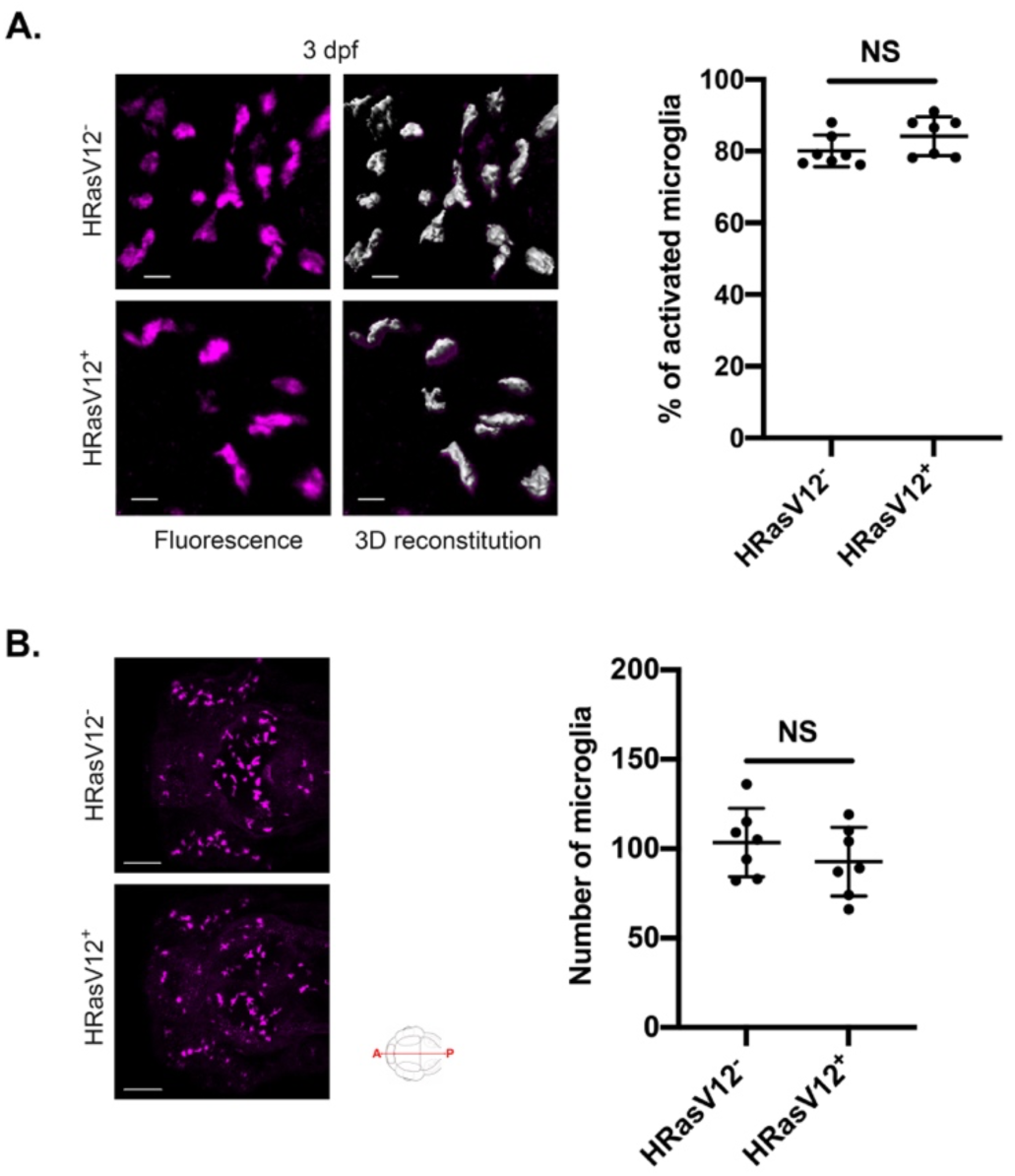
HRasV12 expression in the proliferating domains of the developing CNS doesn’t alter microglia morphology and number at 3 dpf. (A) Close-ups of microglia from 3 dpf HRasV12^-^ (top panels) and HRasV12^+^ (bottom panels) larvae (left panels) and their segmented images in 3D (right panels) using Imaris surface tool, to assess microglia morphology. Scale bar represents 10 μm. The number of activated microglia was quantified within the microglial population of 3 dpf control and HRasV12^+^ larvae. Results are expressed as a percentage of total microglia. HRasV12^-^: n = 7; HRasV12^+^: n = 7; N = 3. Error bars represent mean ± SD. (B) Confocal images of microglia population (magenta) of 3 dpf HRasV12^-^ (top panel) and HRasV12^+^ (lower panel) brains. Scale bar represents 100 μm. The number of microglia from 3 dpf HRasV12^-^ and HRasV12^+^ brains was quantified. HRasV12^-^: n = 7; HRasV12^+^: n = 7; N = 3. Error bars represent mean ± SD. Images were captured using a Zeiss LSM710 confocal microscope with a 20X/NA 0.8 objective. All images represent the maximum intensity projections of Z stacks.

**Sup2:**
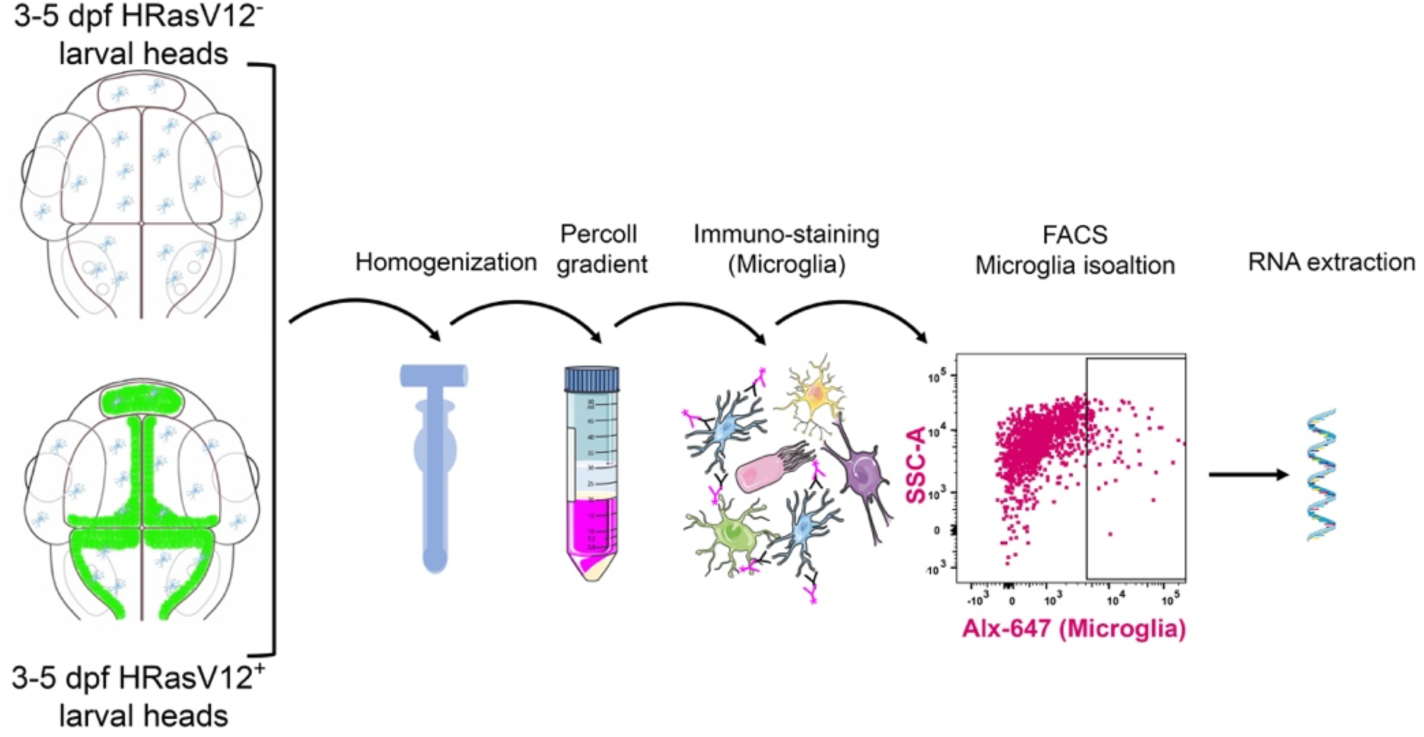
Protocol of microglia isolation from HRasV12^-^ and HRasV12^+^ larvae. Schematic representation of the protocol used to isolate 4C4^+^ microglia from larval zebrafish brains of 3 and 5 dpf HRasV12^-^ and HRasV12^+^ larvae to perform RNA extraction.

**Sup3:**
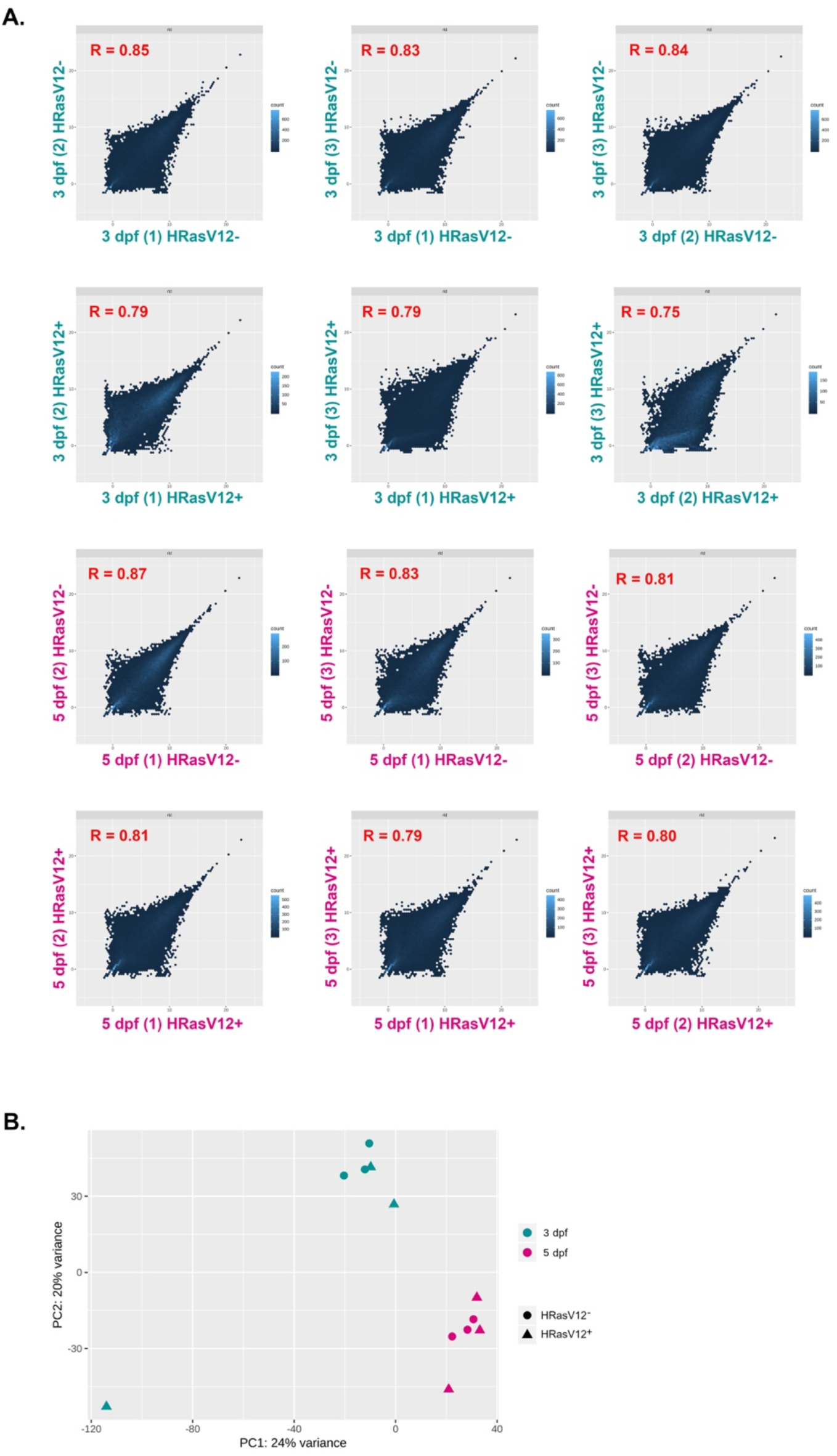
Correlation between biological replicates of isolated microglia from HRasV12^-^ and HRasV12^+^ brains. (A) Normalized counts from 3 dpf replicates 1 and 2, and 1, and 3 HRasV12^-^ and HRasV12^+^ (green). Normalized counts from 5 dpf replicates 1 and 2, and 1, and 3 HRasV12^-^ and HRasV12^+^ (magenta). Pearson’s *r* is indicated. Colours represent point density (Dark blue: low; light blue: high). (B) Principal component analysis (PCA) score plot obtained from normalized counts of isolated microglia from 600 HRasV12^-^ (•) and HRasV12^+^ ( ) larval brains at 3 (green) and 5 (magenta) dpf (N = 3).

**Sup4:**
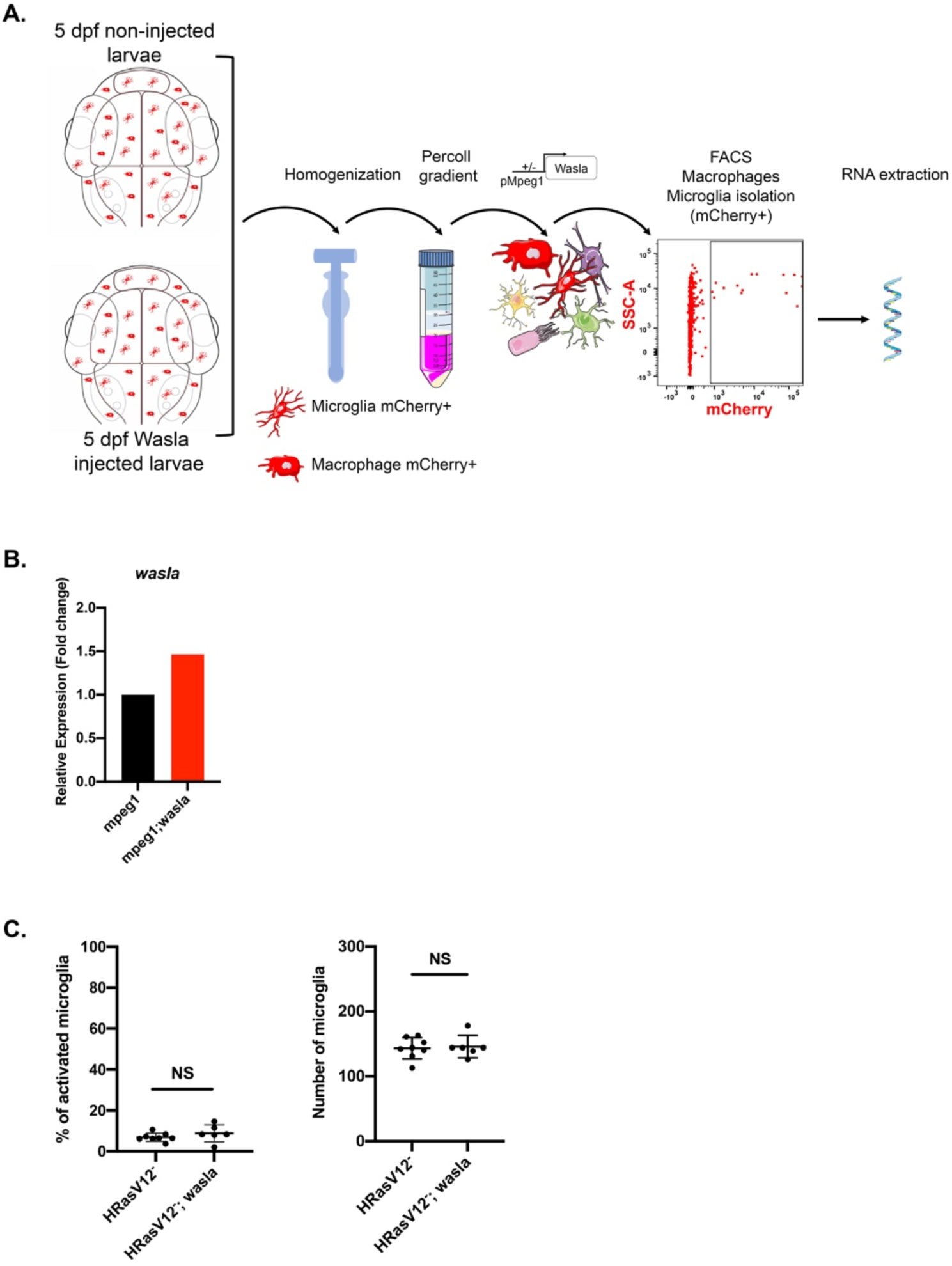
***Walsa* overexpression in microglia from HRasV12^-^ larvae does not affect their number and morphology**. (A) Schematic representation of the protocol used to isolate mCherry+ microglia/macrophages from zebrafish brains of *Tg(mpeg1:mCherry*) 5 dpf larvae non-injected and injected with mpeg1:wasla plasmid to perform RNA extraction. (B) mRNA expression levels of *wasla* from isolated mCherry+ microglia /macrophages from 5 dpf non-injected and injected embryos were determined by qPCR. Fold change is measured in relation to 5 dpf microglia/macrophages using the comparative (ΔΔCT) method. The means are plotted. (C) The percentage of activated microglia and the total number of microglia from 5 dpf HRasV12^-^ and HRasV12^-^;wasla brains were quantified. HRasV12^-^: n = 8; HRasV12^-^; wasla: n = 6; N = 1. Error bars represent mean ± SD

**Tables**

Table1: Actin cytoskeleton genes HRasV12^-^ VS HRasV12^+^

Table2: Phagocytosis of pre-neoplastic cells by FACS

## STAR Methods

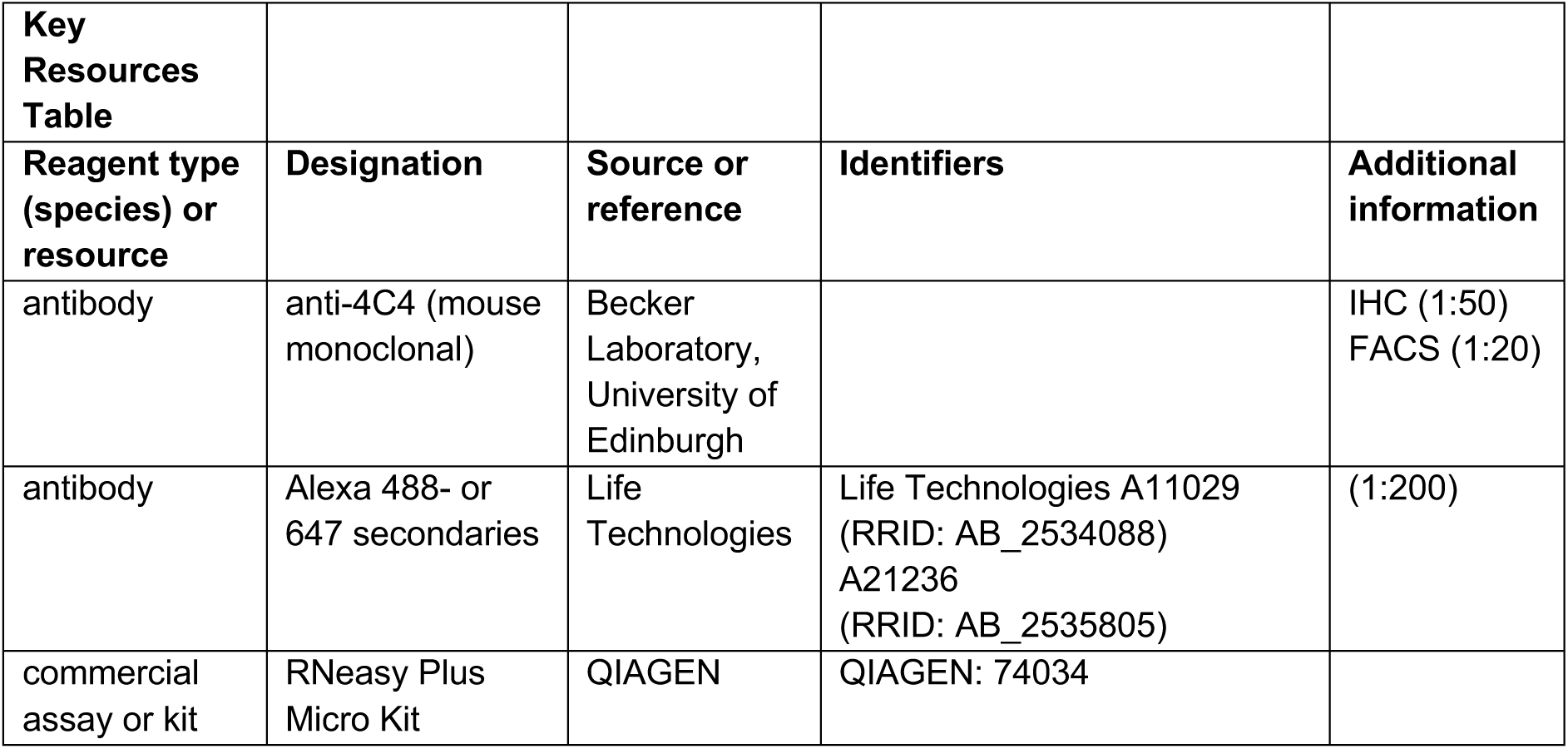
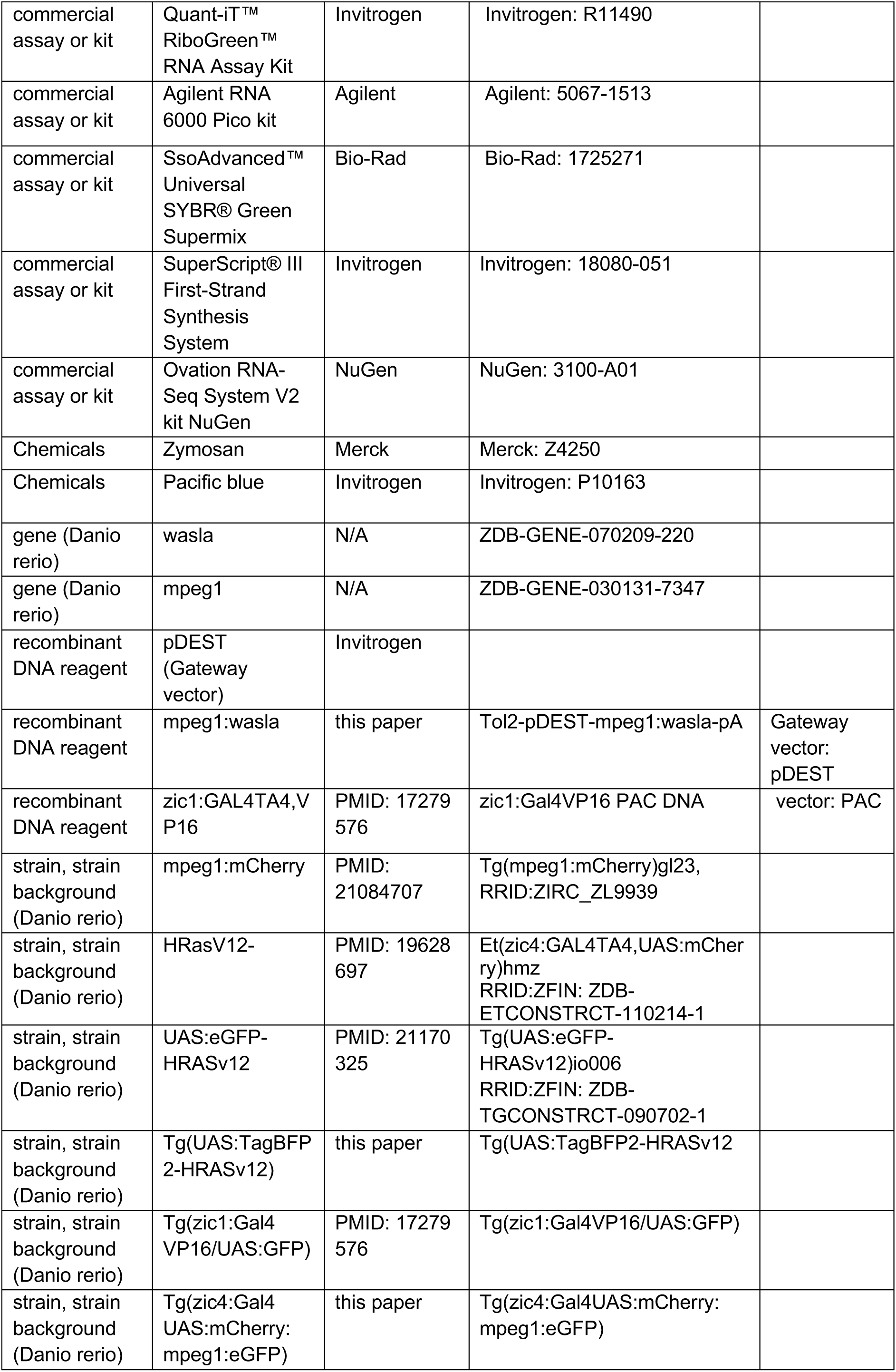
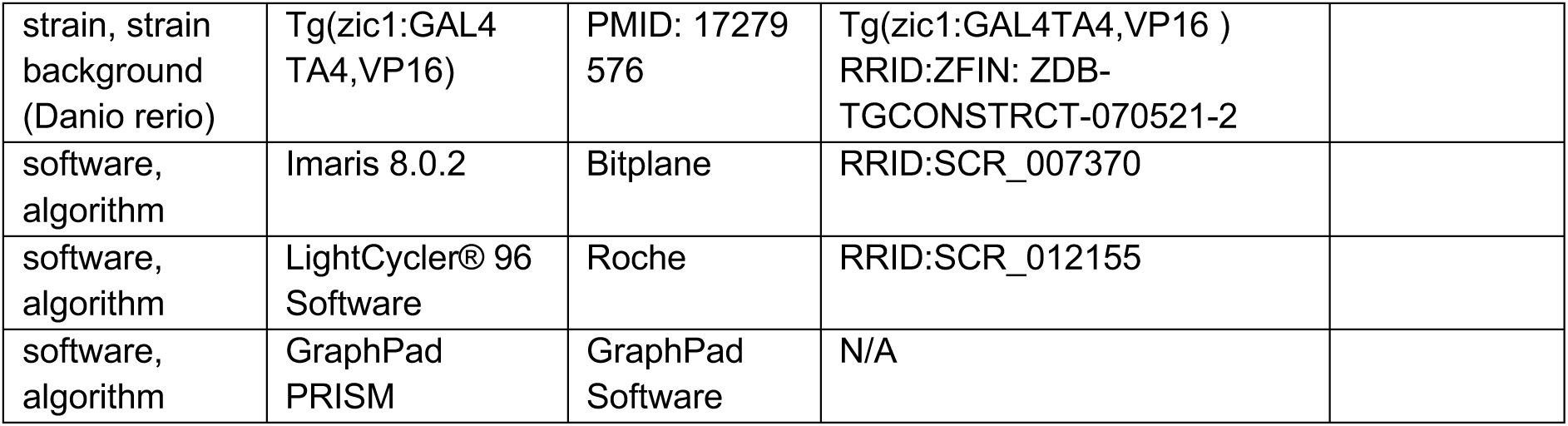

### Zebrafish maintenance

Animal experimentation was approved by the ethical review committee of the University of Edinburgh and the Home Office, in accordance with the Animal (Scientific Procedures) Act 1986. Zebrafish were housed in a purpose-built zebrafish facility, in the Queen’s Medical Research Institute, maintained by the University of Edinburgh Biological Resources. All zebrafish larvae were kept at 28 °C on a 14 hours light/10 hours dark photoperiod. Embryos were obtained by natural spawning from adult *Et(zic4:GAL4TA4,UAS:mCherry)_hmz5_* referred to as HRasV12^-^ (Distel et al., 2009), *Tg(zic1:GAL4TA4,UAS:eGFP)* (Sassa et al., 2007), *Tg(UAS:eGFP-HRasV12)_io006_* (Santoriello et al., 2010), *Tg(mpeg1:mCherry*) (Ellett et al., 2011), *Tg(zic4:Gal4UAS:mCherry:mpeg1:eGFP)*, *Tg(UAS:TagBFP2-HRasV12), Tg(zic1:GAL4TA4,VP16),* and wild-type (WIK) zebrafish strains. Embryos were raised at 28.5°C in embryo medium (E3) and treated with 200 μM 1-phenyl 2-thiourea (PTU) (Sigma) from the end of the first day of development for the duration of the experiment to prevent pigmentation.

### DNA injection to overexpress *wasla*

To generate transient expression of *wasla* in microglia/macrophages of HRasV12^-^, HRasV12^+^ and *Tg(mpeg1:mCherry*) larvae, zebrafish embryos were injected at the one cell stage using an Eppendorf FemtoJet microinjector. Approximately 2 nL of injection solution containing 30 ng/μL of Tol2-pDEST-mpeg1:wasla-pA plasmid, 20 ng/μL Tol2 capped mRNA and 0.1% phenol red were injected into HRasV12^-^, HRasV12^+^ and *Tg(mpeg1:mCherry*) eggs. Larvae were screened at 2 days post-fertilization (dpf) for positive transgene expression and selected for the required experiments.

### Mounting, immunohistochemistry, image acquisition and live imaging

Whole-mount immunostaining of samples was performed as previously described (Astell and Sieger, 2017). Briefly, larvae were fixed in 4% PFA/1% DMSO in PBS at room temperature (RT °C) for 2 hours, then washed in PBStx (0.2% Triton X-100 in 0.01 M PBS) and blocked in 1% goat serum blocking buffer (1% normal goat serum, 1% DMSO, 1% BSA, and 0.7% Triton X-100 in 0.01 M PBS) for 2 hours prior to incubation with the mouse anti-4C4 primary antibody [1:50] overnight at 4 °C to stain microglia. Samples were washed in PBStx before their incubation with conjugated secondary antibodies (goat anti-mouse Alexa Fluor 647 [1:200]) (Life Technologies) overnight at 4 °C. The samples were washed several times with PBStx and stored in 70% glycerol at 4 °C until final mounting in 1.5% low melting point agarose (Life Technologies) in E3 for image acquisition. Whole-brain immunofluorescent images were acquired using confocal laser scanning microscopy (Zeiss LSM710 and LSM880; 20× objective (air, NA = 0.8); z-step = 2.30 μm; 405 nm, 488 nm, 594 nm, 633 nm laser lines).

Live imaging of zebrafish larvae was performed as previously described (Chia et al., 2018); samples were anaesthetized with 450 μM Tricaine (MS222, Sigma) and mounted dorsal side up in 1.5% low melting point agarose (Life Technologies), in 60 x 15 mm petri dishes (Corning) filled with E3 containing 450 μM Tricaine and 200 μM PTU. To investigate microglia motility and phagocytosis from 3 and 5 dpf HRasV12^-^, HRasV12^+^ and HRasV12^+^/wasla larval brains, time-lapse imaging with Z stacks were acquired using a Zeiss LSM880 confocal microscope equipped with an Airyscan Fast module and a piezo z-drive (Zeiss Plan-Apochromat 20× (water, NA = 1.0); z-step = 1.5 μm). All time-lapse acquisitions were carried out in temperature-controlled climate chambers set to 28 °C for 13 hours with acquisition every 14 min.

### Proliferation assay (EdU staining)

Proliferation assay was performed following guidance of Click-iT™ EdU Cell Proliferation Kit for Imaging, Alexa Fluor™ 647 dye (Invitrogen). Briefly, embryos were collected from the outcross of the driver fish line *Tg(zic4:GAL4TA4,VP16)* to the *Tg(UAS:TagBFP2-HRasV12)* fish lines. At 2 dpf dechorionated HRasV12^-^ and HRasV12^+^ larvae were incubated with EdU (50 μM) in E3 containing 200 μM PTU then raised at 28.5 °C. At 5 dpf larvae were fixed in 4% PFA/1%DMSO in PBS at RT °C for 2 hours, washed several times in PBStx, digested with collagenase (Sigma) at 2 mg/ml in PBS at RT °C for 40 min under agitation, washed in PBStx then incubated in Click-iT reaction cocktail at RT °C for 4 hours under agitation. Larvae were washed in PBStx then blocked in 1% goat serum blocking buffer to carry out microglia immunostaining as described in previous paragraph.

### Phagocytosis assay

Phagocytosis assay was performed using an Eppendorf FemtoJet microinjector to inject custom made zymosan (Sigma) coupled with Pacific Blue fluorochrome (molecular probes) into the larval zebrafish brain. Anesthetised 3 and 5 dpf HRasV12^-^, HRasV12^+^ and HRasV12^+^/wasla larvae were injected into either telencephalon or tectum with approximately 2 nL of injection solution composed of 2.5.10^5^ zymosan/μL, 0.1% phenol red in PBS. Injected larvae were maintained at 28.5 °C in E3 containing 200 μM PTU for 6 hours, fixed in 4% PFA/1% DMSO in PBS at RT °C for 2 hours then microglia immunostaining was performed.

### Image analysing

Analysis of all images was performed in 3D using Imaris (Bitplane, Zurich, Switzerland). To assess microglia (4C4^+^) morphology and volume measurements, we used the surface-rendering tool in Imaris 8.2.1, which allowed segmentation of individual cells in 3D as well as bigger volumes such as brain and pre-neoplastic mass volume. To assess microglia morphological activation, we calculated the ratio of the cellular surface and cellular volume of individual cells as previously described (Chia et al., 2018; Gyoneva et al., 2014). Microglia with a ratio smaller than 0.8 were classified as activated. The ‘Spots’ function tool was used to quantify the number of activated microglia related to the total number of microglia within the full brain, and the averaged value expressed as measure of the percentage of activated microglia. To quantify proliferation rates, the number of 4C4^+^/EdU^+^ cells (EdU^+^ microglia) were counted in relation to the total number of microglia within the full brain and the averaged value expressed as measure of percentage of microglia proliferation. To quantify zymosan phagocytosed by microglia of 3 and 5 dpf HRasV12^-^, HRasV12^+^ and HRasV12^+^/wasla larval brains, we used the surface-rendering tool in Imaris 8.2.1 to manually create a surface corresponding to the fluorescent signal of all zymosan particles. An additional surface for microglia allowed us to distinguish zymosan that had been phagocytosed by microglia. This allowed us to read out the sum of fluorescence intensity (AU) of phagocytosed zymosan within microglia surfaces and the total AU of all zymosan surfaces. The percentage of microglia phagocytosis was calculated from the averaged value of the number of phagocytosed zymosan related to the total number of zymosan within either the telencephalon or the tectum (Engulfed zymosan fluorescent intensity:total zymosan fluorescent intensity) x 100).

To assess microglia motility of 3 and 5 dpf HRasV12^-^, HRasV12^+^ and HRasV12^+^/wasla larval brains, we used the ‘Tracking’ function tool and manually tracked mpeg:eGFP^+^ cells along time. Imaris software calculated track length and track speed mean for each cell and microglia tracks were displayed as time color-coded lines for the different conditions. To measure the number of phagosomes per microglia from 5 dpf HRasV12^+^ and HRasV12^+^/wasla larvae, we used mpeg1:eGFP^+^ cells to visualise phagosomes in black within GFP^+^ cytoplasm. The combination of 3D and slide views on Imaris software allowed us to manually count the number of phagosomes in microglia within the tectum of 10 larvae per condition.

### Microglia/macrophage isolation and FACS analysis

Microglia were isolated by FACS from 3 and 5 dpf heads of HRasV12^-^ and HRasV12^+^ larvae as previously described (Mazzolini et al., 2018) whereas microglia and macrophages were isolated from whole 5 dpf mpeg1:mCherry^+^ and mpeg1:mCherry^+^/wasla larvae. FACS allowed cell separation from debris in function of their size (FSC-A) and granularity (SSC-A). Single cells were then separated from doublets or cell agglomerates (FSC Singlet; SSC Singlet). From the single-cell population, a gate was drawn to separate live cells (DAPI−) from dead cells (DAPI+). Unstained and cells incubated with secondary antibody Alx647 only were used as controls to draw gates corresponding to microglia (4C4^+^/Alx647^+^) populations. Finally, microglia (4C4^+^/Alx647^+^; Sup2) and microglia/macrophage (mCherry^+^; Sup4A) were segregated from the live cell population gates. FACS data were analyzed using FlowJo Software (Treestar, Ashland, OR).

### RNA extraction and cDNA amplification

All experiments were performed in three replicates with a total number of 600 larvae per replicate. Total RNA extraction from microglial cells was performed using the Qiagen RNeasy Plus Micro kit according to the manufacturer’s guidance (Qiagen). RNA sample quality and concentration were determined using Agilent RNA 6000 Pico kit and a Agilent 2100 Bioanalyser System (Agilent Technologies).

For sequencing, all RNA samples with a RIN score > 7 were transcribed into cDNA using the Ovation RNA-Seq System V2 kit according to the manufacturer’s instructions (NuGEN). Samples were then sent to Edinburgh Genomics for library synthesis and sequencing. For qPCR, RNA sample quality and concentration were assessed using the LabChip GX Touch Nucleic Acid Analyzer and RNA Pico Sensitivity Assay. All RNA samples with a RIN score > 7 were transcribed from the same amount of RNA into cDNA using the Super-Script® III First-Strand Synthesis System (Invitrogen).

### Library synthesis

Sequencing libraries were prepared using the Illumina TruSeq DNA Nano library preparation kit according to manufacturer’s instructions with amended shearing conditions (duty factor 10%, PIP 175, cycles/burst 200, duration 40 s) using a 500 ng input of amplified cDNA (Illumina, Inc.). The size selection for the sheared cDNA was set for 350 bp products. Libraries were normalized and ran on 2 HiSeq 4000 lanes with 75-base paired-end reads resulting in an average read depth of around 20 million read pairs per sample.

### Bioinformatics

The quality control of the sequences was done with FastQC (Andrews, 2010), and Trimmomatic was applied to trim low-quality reads and adapters (Bolger et al., 2014). We aligned the RNA-seq reads to the zebrafish reference genome (Ensembl, GRCz11) using STAR v2.6 (Dobin et al., 2013) and transcript were assembled and counted with HTSeq (Anders et al., 2015) using annotation from Ensembl (Danio_rerio.GRCz11.93.gtf). Counts normalization, transformation (rlog), and differential expression analysis were performed using DESeq2 (Love et al., 2014). Normalized data were inspected using Principal Component Analysis (PCA) (Sup3B), and inter-sample correlation plots (Sup3A). We selected genes related to actin cytoskeleton among the KEGG database: dre04810 “Regulation of actin cytoskeleton”, dre04510 “Focal adhesion”, dre04520 “Adherens junction”, dre04150 “mTOR signaling pathway”, and dre04530 “Tight junction” (PMID: 10592173). We performed the gene expression comparison between isolated microglia from 3 dpf and 5 dpf HRasV12^-^ and HRasV12^+^ larvae using DESeq2. Finally, we looked for top ranked genes differentially expressed at 5 dpf and constant at 3 dpf in HRasV12^-^ and HRasV12^+^ microglia, for further investigations.

### Quantitative PCR

Quantitative (qPCR) amplifications were performed in technical triplicates in a 20 μl reaction volume containing SsoAdvanced Universal SYBR Green Supermix (Bio-Rad) using a LightCycler 96 Real-Time PCR System (Roche). The PCR protocol used was initial denaturation step of 10 min at 95 °C, and 45 cycles of 10 s at 95 °C, 20 s at 56 °C, and 20 s at 72 °C. Primers used were:

Beta-actin forward 5’-CACTGAGGCTCCCCTGAATCCC-3’.

Beta-actin reverse 5’-CGTACAGAGAGAGCACAGCCTGG-3’.

wasla forward 5’-CAAATGTGGGCTCTCTCCTTCTGAC-3’.

wasla reverse 5’-GAGGGTTGTCTTTCACCAAACAGGC-3’.

Melting curve analysis was used to ensure primer specificity. For qPCR analysis, the threshold cycle (Ct) values for each gene were normalized to expression levels of ß- actin and relative quantification of gene expression determined with the comparative Ct (ΔΔCt) method using the LightCycler® 96 Software (Roche).

### Survival assay

HRasV12^-^, HRasV12^+^ and HRasV12^+^/wasla larvae were screened at 2 dpf for positive transgene expression then were housed in a purpose-built zebrafish facility, in the Queen’s Medical Research Institute, maintained by the University of Edinburgh Biological Resources. Larvae were kept by 20 per nursing tanks (3 replicates) at 28 °C on a 14 hours light/10 hours dark photoperiod, daily fed by facility’s staff members from 5 dpf to 31 dpf. Surviving larvae were counted every day for 1 mpf.

### Statistical analysis

All experiments were performed in three replicates. All measured data were analysed using StatPlus (AnalystSoft Inc.). Unpaired two-tailed Student’s t-tests were performed to compare two experimental groups, and one-way ANOVA with Bonferroni’s post-hoc tests for comparisons between multiple experimental groups. Statistical values of p < 0.05 were considered to be significant. All graphs were plotted in Prism 8 (GraphPad Software) and values presented as population means ± SD.

